# Moving from Association to Causation: Instrumental factor models for causal inference in high-dimensional multi-omics data

**DOI:** 10.64898/2025.12.26.696613

**Authors:** Aditya Mishra, Michelle Badri, Elizabeth Coler, Zhe Liu, Richard Bonneau, James T. Morton, Alexander A. Aksenov

## Abstract

Distinguishing true causal relationships from mere association is a core challenge in bioinformatics and science in general. Causal inference, in particular in “-omics” studies, is hindered by high dimensionality, correlated measurements, and pervasive endogeneity arising from unmeasured confounding and complex interventions. We introduce Factor IV, a supervised instrumental variable (IV) framework for identifying causal effects in high-dimensional biological systems. FactorIV constructs low-dimensional instrumental factors via a sparse low-rank decomposition of the instrument-exposure map, yielding an identifiable first stage even when either/both endogenous and instrumental variables are high-dimensional. Under standard IV assumptions, these factors remain orthogonal to outcome noise while capturing coordinated, perturbation-driven biological variation. The framework supports generalized first-stage models, including Gaussian, Bernoulli and negative binomial likelihoods. Simulation studies demonstrate accurate recovery of factor-level and feature-level causal effects under linear and generalized settings, with robustness to correlated errors and hidden confounding. To illustrate biological discovery, we applied FactorIV to a mouse hepatocellular carcinoma model and the DIABIMMUNE infant cohort. We demonstrate how FactorIV uncovers mechanistically interpretable causal structure beyond association-based analyses.

## 1 Main

The ultimate goal of biology is to identify causal pathways whose perturbation can reliably alter biological and clinical outcomes. This objective lies at the core of precision medicine but remains difficult to achieve because biological systems involve multiple interacting pathways and many unmeasured factors. While statistical associations are easy to detect, they rarely distinguish causal drivers from observational data. Traditional approaches rely on molecular validation through cultivation [1] and genetic manipulation [2] within randomized experimental designs [3]. However, these strategies are limited because only a small fraction of microbes are culturable [4], and microbial function often emerges from complex ecological interactions that cannot be reduced to single-variable relationships [5].

Similar challenges arise in metabolomics and other “omics” layers, where metabolites frequently participate in multiple pathways and cross-feeding interactions. A given compound may be synthesized by one organism, modified by another, or reflect downstream signaling cascades across the community. These multi-layered dependencies generate dense causal networks that are difficult to resolve without appropriate statistical frameworks.

Advances in high-throughput sequencing have dramatically expanded our knowledge of uncultured microorganisms [6–9], revealing extensive links between the microbiome, diet, host metabolism, and immune phenotypes, and highlighting opportunities for clinical intervention [10, 11]. However, most microbiome studies cannot control for all biological and technical confounders due to environmental variability, cohort constraints, and the infeasibility of experimentally manipulating complex microbial ecosystems. As a result, observational analyses often struggle to uncover causal mechanisms underlying key organisms and molecular biomarkers.

Experimental designs that apply structured interventions, such as genetic knockouts, antibiotic exposure, or dietary shifts, offer opportunities to infer causal relationships between microbes and host phenotypes. Instrumental variable (IV) methods provide a principled framework for leveraging such designs [12, 13]. In this setting, a set of instruments **Z** ∈ ℝ^*n×d*^ influences the microbial community **X** ∈ ℝ^*n×p*^ but affects the outcome **Y** ∈ ℝ^*n×q*^ only through **X** and is independent of both observed and unobserved confounders (Figure 1A). Observed confounders **Q** ∈ ℝ^*n×t*^ may include age, sex, or batch effects, while unobserved confounders **H** ∈ ℝ^*n×s*^ may arise from uncontrolled diet, environment, or sample processing. IV methods aim to isolate exogenous variation in **X** that is not driven by these factors.

**Fig. 1.**
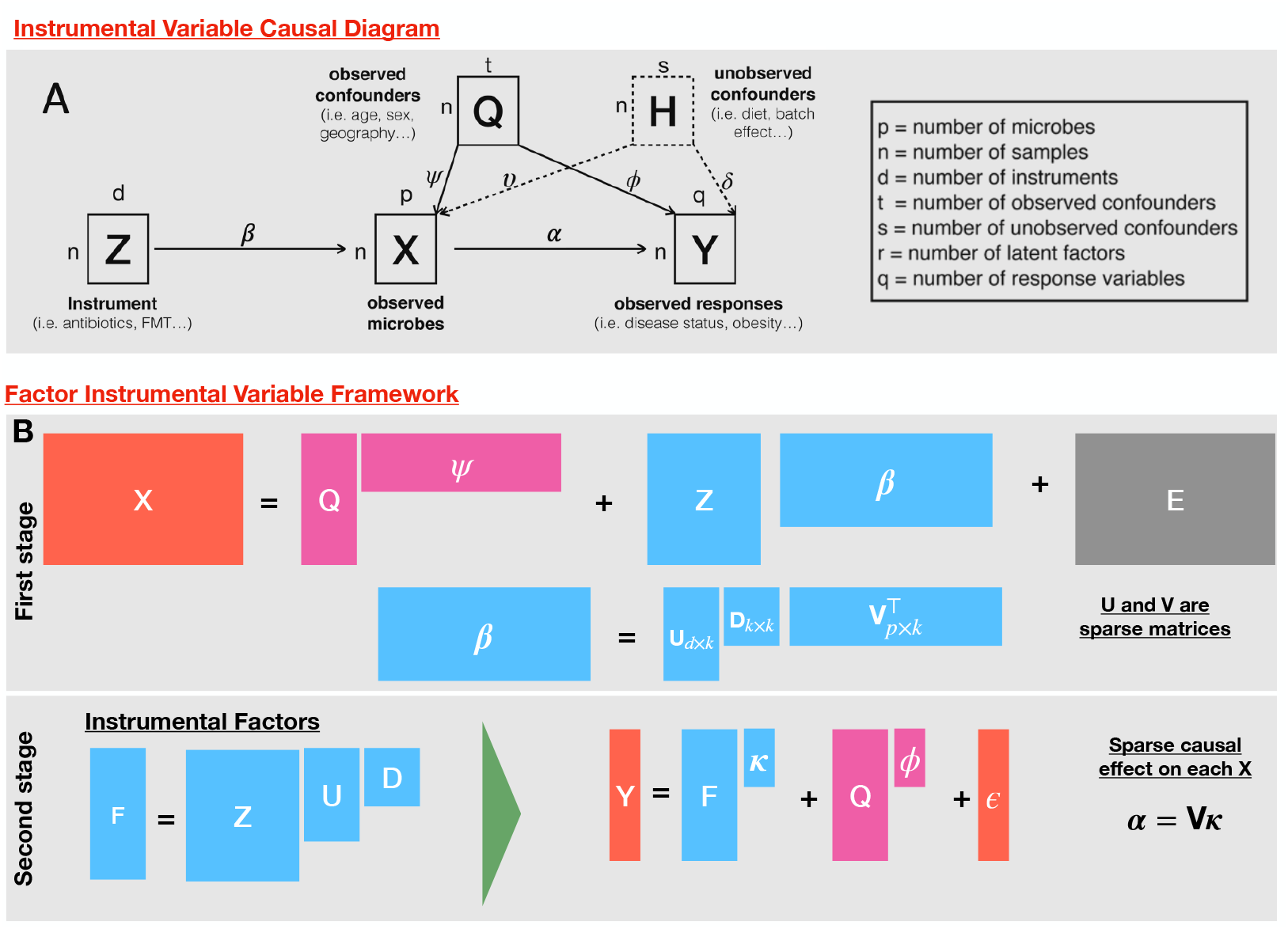
Illustration of instrumental variable analysis and the Factor IV framework. (A) Instrumental variable causal diagram, where **Z** denotes experimental perturbations, **X** observed microbial features, **Y** outcomes, **Q** observed confounders, and **H** unobserved confounders affecting both **X** and **Y**. (B) Factor IV replaces direct instrument usage with a supervised low-rank factorization of the instrument–exposure map, yielding instrumental factors **F** = **ZUD** that summarize perturbation-driven variation in **X**. Causal effects are estimated at the factor level via ***κ*** and mapped to individual features through ***α*** = **V*κ***.

Despite their conceptual appeal, applying IV methods to multi-omics data presents substantial statistical challenges. Classical IV analysis requires sufficient instrument variation relative to model complexity and becomes unstable when the number of molecular features far exceeds the sample size (*n* ≪ *p*) [12, 14]. Related approaches such as Mendelian randomization incorporate regularization [15, 16], but are not directly suited to microbiome data, which are sparse, zero-inflated, and compositional [17, 18]. In these settings, non-causal correlations arise not only from confounding but also from the relative abundance constraint that couples all components of **X**, further complicating identifiability and interpretation.

To address these challenges, we develop a sparse Factor IV framework (Figure 1B) that combines supervised factor analysis with instrumental variables [19–22]. By projecting high-dimensional multi-omics measurements onto a low-dimensional set of latent factors anchored by experimental instruments, Factor IV stabilizes estimation in the *n* ≪ *p* regime while preserving biologically meaningful structure. The framework accommodates count and compositional data through appropriate likelihoods, including Poisson and negative binomial models, and accounts for unobserved confounding through instrument-aligned latent structure. Through simulations and two biological case studies—autoimmune disease risk in infants predisposed to type I diabetes and a mouse model of liver cancer—we show that Factor IV both recapitulates known causal relationships and uncovers additional biologically interpretable patterns that are not detectable using standard association-based methods such as DESeq2 [23].

## 2 Results

### 2.1 Simulation benchmarks demonstrate accurate recovery of causal effects and robustness to model misspecification

We first evaluated Factor IV under controlled simulation settings where the true data-generating process is known. To reflect the complexity of modern multi-omics experiments, we simulated datasets with high-dimensional endogenous variables (*p* ≫ *n*), correlated feature structure, sparse causal effects, and multiple sources of endogeneity. The simulations included both linear and generalized linear first-stage relationships, mimicking continuous and count-based omics data. In this regime, classical instrumental variable estimators are not identifiable because the first-stage regression becomes ill-posed when the number of features exceeds the sample size [24]. Factor IV addresses this instability by imposing a supervised low-rank structure on the instrument–feature map, yielding an identifiable first stage even under high dimensionality.

Data were generated from a latent-factor IV model (Fig. 1) with *k* latent biological factors, *d* instruments, *p* endogenous features, and *t* observed confounders, with an additional subset of features affected by unobserved confounders **H**. The endogenous matrix **X** followed a structured low-rank model in which instruments **Z** act through a sparse coefficient matrix ***β*** = **UDV**^⊤^, producing interpretable perturbation patterns consistent with realistic multi-omics experiments. Outcomes **Y** were generated from the latent factors via a sparse structured effect ***α*** = **V*κ***, together with contributions from both observed and unobserved confounders (Fig. 1A).

Endogeneity was introduced through two biologically motivated mechanisms: (i) unobserved confounders influencing both stages, and (ii) correlated first- and second-stage error terms inducing dependence between **X** and **Y**. These mechanisms are formally equivalent under the observed-data model and replicate common biological failure modes in which upstream processes (e.g., diet, inflammation, or batch effects) jointly affect molecular features and host phenotypes (Methods; Supplementary Section S8). Signal-to-noise ratios for both **X** and **Y** were explicitly controlled to span realistic effect sizes.

Under correct model specification, Factor IV accurately recovered all ground-truth quantities across both Gaussian and negative binomial settings (see Figure 2). Instrumental factors were estimated using GOFAR for continuous data and NBFAR for count data, yielding low-dimensional components 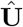 and loadings 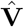 that capture instrument-aligned variation [19, 20]. In the second stage, outcomes were regressed on the estimated instrumental factors 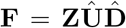 to obtain 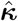, and feature-level causal effects were reconstructed as 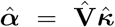. Factor IV nearly perfectly reconstructed the outcome **Y**, factor-level effects ***κ***, and instrumental signal 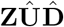, demonstrating that supervised factorization resolves the *p* ≫ *n* identifiability barrier and enables consistent causal estimation under substantial noise.

**Fig. 2.**
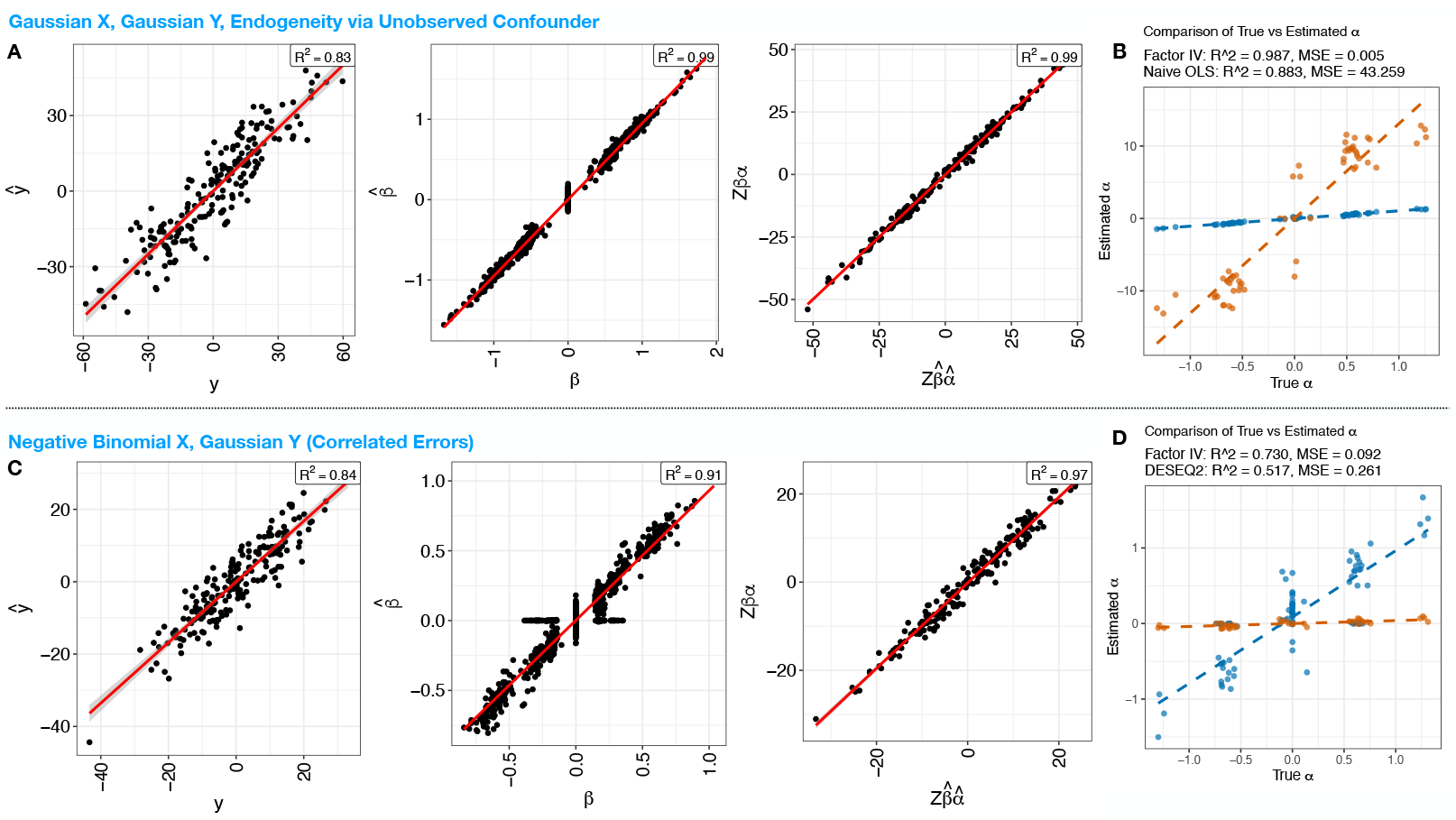
Simulation benchmarks for the Factor IV framework. Performance under Gaussian and negative binomial exposure models in the presence of endogeneity induced by unobserved confounding and correlated errors. (**A**) Gaussian setting: observed versus predicted outcomes **Y**, instrument–exposure signal **Z*β***, and projected causal component 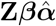. (**B**) Gaussian setting: comparison of estimated feature-level causal effects 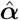 from Factor IV and naive regression. (**C**) Negative binomial setting: observed versus predicted outcomes **Y**, instrument–exposure signal **Z*β***, and projected causal component 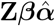. (**D**) Negative binomial setting: comparison of estimated causal effects 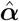 from Factor IV and DESeq2.

Figure 2A,C (and Supplementary Fig. 1A,C) summarizes performance across all four simulation scenarios. Factor IV consistently recovered the true causal effects in both linear and generalized linear settings, accurately estimating the instrument– exposure map 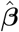 and reconstructing the projected causal signal 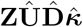 [22, 25]. These results confirm that the supervised factor stage isolates exogenous variation in **X** and that the second-stage regression recovers the correct causal effects.

We compared Factor IV with commonly used association-based methods that implicitly assume the reverse causal direction (**Y** → **X**) (see Figure 2B,D (and Supplementary Fig. 1)B,D). For continuous features, we applied feature-wise ordinary least squares regression, and for count data we used DESeq2 [23]. In both cases, association-based approaches produced numerous false positives driven by endogeneity and correlated noise. In contrast, Factor IV consistently recovered the true causal effects, highlighting the limitations of association-based analyses for causal interpretation and the necessity of high-dimensional IV frameworks in multi-omics studies.

### 2.2 Effect of early-life microbiome on infant development

After validating Factor IV using synthetic benchmarks, we applied the framework to the DIABIMMUNE infant cohort to characterize age-specific microbial contributions to early immune development. The results are summarized in Fig. 3. Consistent with prior analyses [26], Factor IV recapitulates known associations between the infant gut microbiome and autoimmune risk, while additionally resolving temporal patterns that are not detectable using conventional taxonomic profiling.

**Fig. 3.**
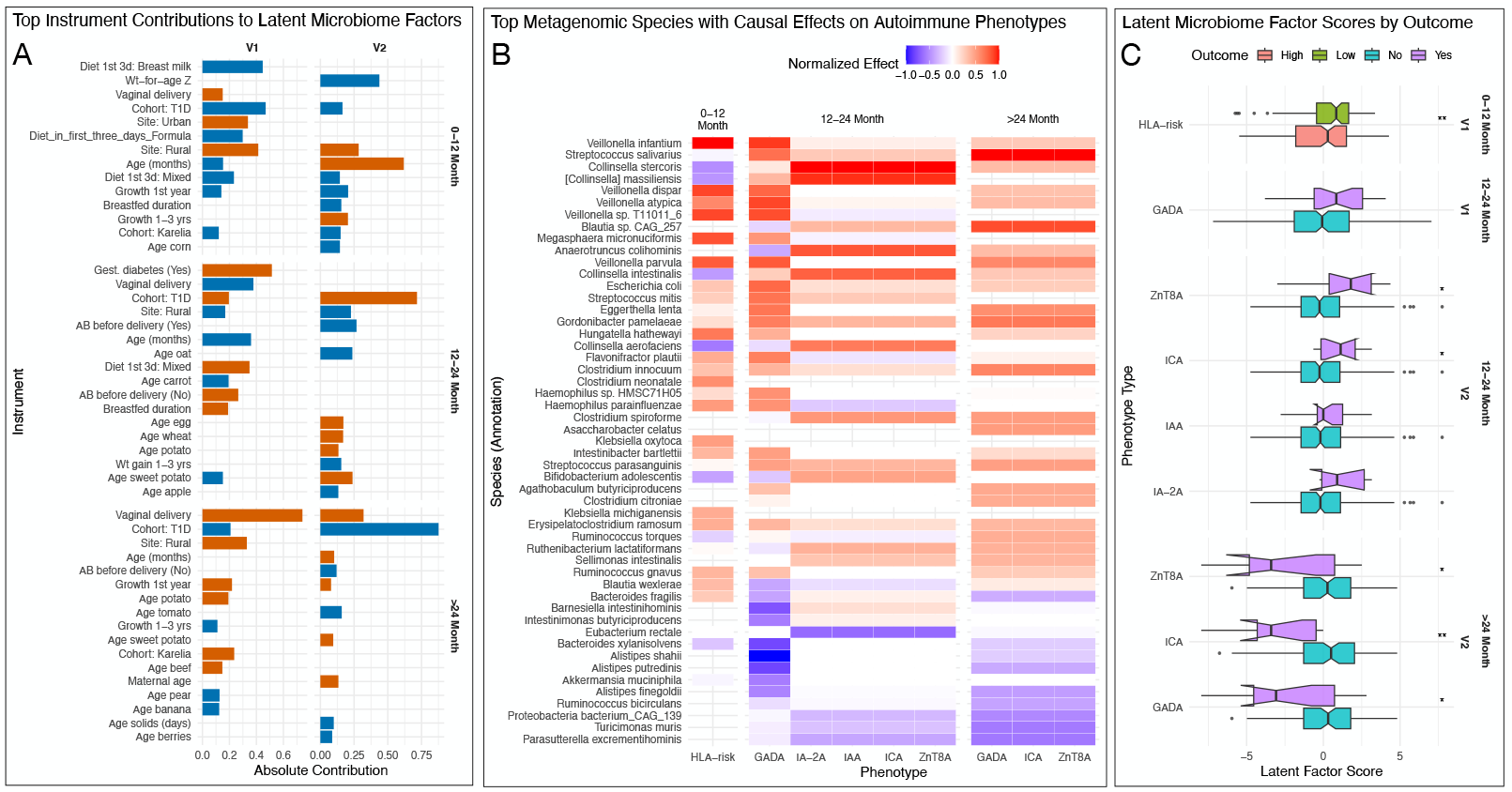
Age-stratified Factor IV analysis of the infant gut microbiome. Latent microbiome factors integrating metagenomic profiles, early-life instruments, and autoimmune phenotypes across developmental windows. (**A**) Contributions of early-life instruments (diet, antibiotic exposure, delivery mode, and environmental factors) to each instrumental factor, estimated separately by age group; bar length indicates magnitude and color indicates direction. (**B**) Heatmap of normalized feature-level causal effects of metagenomic species on autoimmune phenotypes across three age strata (0–12 months, 12–24 months, and *>* 24 months), showing taxa with the largest absolute effects. (**C**) Box plots of instrumental factor scores stratified by autoimmune outcome group, illustrating age-specific associations between perturbation-aligned microbial structure and host phenotypes; notches indicate median confidence intervals, with significance assessed using two-sided tests (p *<* 0.05).

Fig. 3A shows the relative contributions of genetic, environmental, and clinical instruments to latent microbiome factors across three developmental windows. During the first year of life (0–12 months), HLA risk status dominates microbiome structure, whereas delivery mode, breastfeeding, and environmental exposures become increasingly influential thereafter, consistent with established drivers of microbiome maturation [27]. This age stratification reveals a transition from genetically driven to environmentally driven microbiome organization.

Evidence supporting the original findings are shown in Panel B, where *Ruminococcus gnavus* shows strong positive associations with autoimmune phenotypes, particularly during early infancy. This pathobiont produces inflammatory glucorhamnan polysaccharides that induce TNF*α* secretion via TLR4 signaling [28], providing a mechanistic explanation for its role in the *α*-diversity decline preceding T1D onset [26]. Multiple Streptococcus species also show positive correlations with autoimmune phenotypes, corroborating the original identification of inflammation-associated organisms during the critical seroconversion window.

Beyond these known signals, Factor IV identifies taxa with protective effects that are not evident in association-based analyses. Notably, *Alistipes shahii* and *Akkermansia muciniphila* exhibit consistent inverse effects on autoimmune phenotypes across all developmental windows. The protective role of *A. muciniphila* is supported by experimental evidence demonstrating its ability to enhance intestinal barrier integrity, reduce endotoxemia, and promote regulatory T cell differentiation [29, 30]. Recent studies in NOD mice further show that *A. muciniphila* supplementation delays T1D onset by inducing tolerogenic dendritic cells and shifting immune balance toward Treg populations [31].

The temporal specificity revealed in Panel Fig. 3C represents an important advancement in understanding of T1D pathogenesis. The predominance of HLA-risk associations exclusively in the 0-12 month period suggests a critical window where genetic predisposition most strongly influences microbiome assembly and immune development. This observation supports the “critical period hypothesis” and indicates that therapeutic interventions may be most effective during early infancy when genetic and environmental factors intersect.

The age-dependent shifts in microbial effects revealed by the factor analysis are consistent with the evolving functional requirements of the developing immune system. During the 0-12 month period, when adaptive immunity is still maturing, inflammatory species like *R. gnavus* may disproportionately influence immune programming through innate pathways. The increasing prominence of protective species like *A. muciniphila* in later periods suggests that as immune tolerance mechanisms mature, anti-inflammatory microbes become more important for maintaining homeostasis. From a mechanistic perspective, these findings suggest pathways where early inflammatory damage from pathobionts like *R. gnavus* compromise intestinal barrier integrity [32], facilitating translocation of microbial antigens that trigger autoimmune cascades. Simultaneously, lack of protective species like *A. muciniphila* reduces the microbiome’s capacity to maintain immune tolerance through SCFA production and regulatory T cell induction [33].

The Factor IV analysis approach uncovers latent microbiome-host interactions patterns across multiple data dimensions - taxonomic, functional, and temporal. These uncovered age-specific therapeutic windows and distinct microbial contributions could potentially provide a framework for developing microbiome interventions in T1D mitigation. Future studies should investigate whether early-life supplementation with protective species like *A. muciniphila*, combined with targeted reduction of pathobionts during critical developmental windows, can delay T1D onset in high-risk populations.

### 2.3 Multi-omics analysis of a murine HCC model

We applied Factor IV to a murine hepatocellular carcinoma (HCC) study integrating gut microbiome, serum metabolomics, immune perturbations, and dietary interventions [34]. The experimental design included multiple, overlapping perturbations—high-fat diet (HFD), antibiotics, and immune-related genetic knock-outs—making it well suited for instrumental variable analysis in a high-dimensional setting.

Diet, antibiotics, and genetic knockouts were treated as instruments, with age, strain, and body weight included as observed confounders. Microbiome and metabolome profiles were modeled jointly in the first stage using generalized co-sparse factor regression (GOFAR) [19, 35], yielding two instrumental factors with corresponding loadings (**V1** and **V2**) that captured the dominant instrument-induced variation (Fig. 4). These factors were then used in the second-stage regression to estimate causal effects on multiple host phenotypes, including obesity, NASH, IgA status, B-cell depletion, and HCC.

**Fig. 4.**
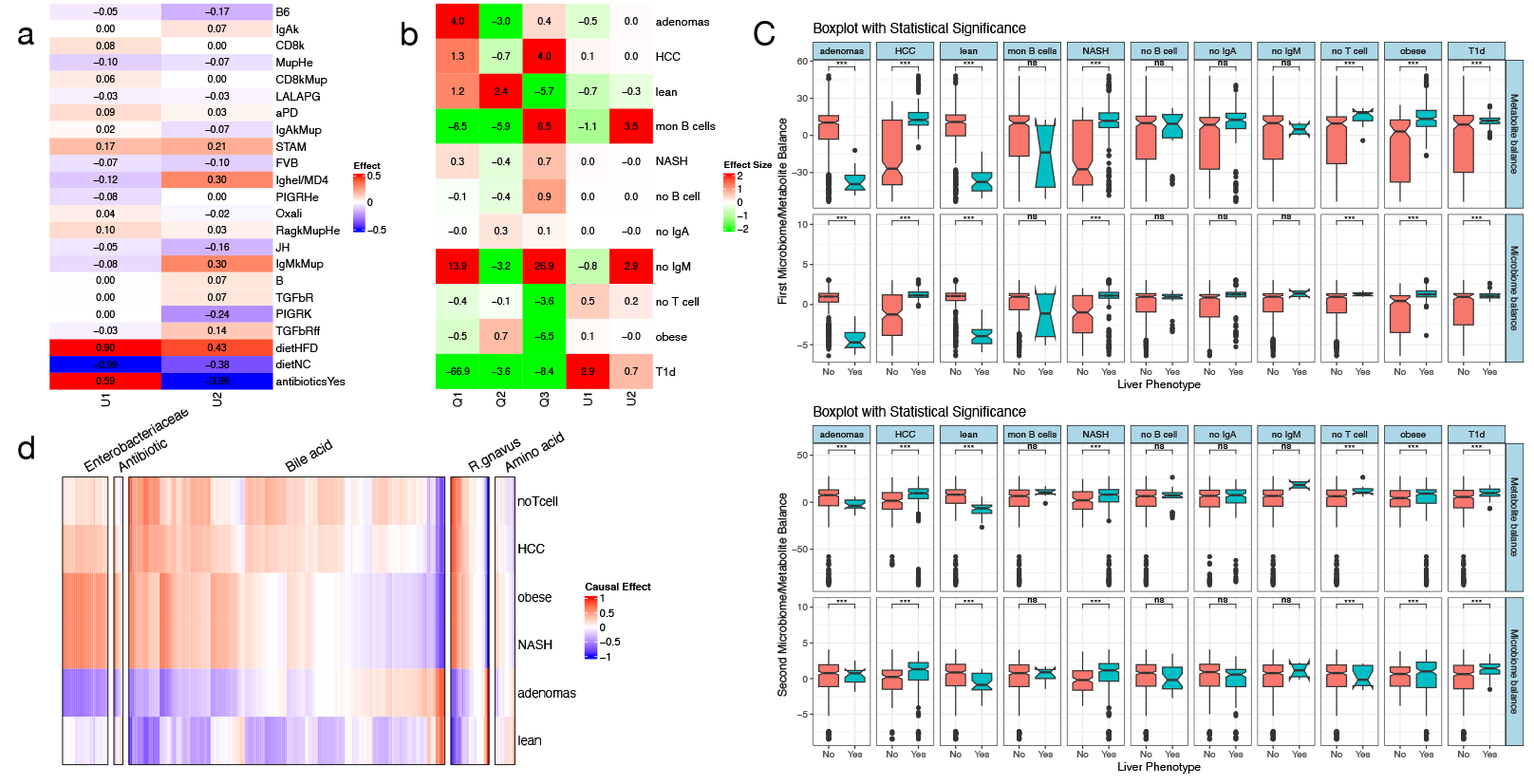
Application of the FactorIV approach to analyze the causal effects of microbiome and metabolite profiles on the occurrence of various stages of liver inflammation. (a) Relative importance of genetic mutations and dietary interventions in forming two latent factors that represent underlying biological processes. (b) Causal effects of the latent factors, alongside observed confounders, on the progression through various stages of liver inflammation. (c) Relative contributions of microbiome and metabolite balances, as identified by loadings **V**_1_ and **V**_2_, in distinguishing between the stages of liver inflammation. (d) Specific causal effects of key microbiome species and metabolite markers on the progression of liver inflammation across its various stages.

Inspection of the estimated instrument loadings (**U**) showed that HFD and antibiotics were the primary drivers of the learned factors (Fig. 4A), consistent with the original experimental findings [34]. Additional contributions from knockouts such as *IgA*^*k*^*Mup, Mup*^*He*^, STAM, and B6 reflected their more moderate but biologically interpretable effects on the microbiome and metabolome. Among the two factors, only **V1** was significantly associated with obesity, NASH, and HCC (Fig. 4C), whereas **V2** showed weak or no association with disease outcomes. As a result, estimation of feature-level causal effects ***α*** = **V*κ*** was effectively driven by **V1**, with ***α*** reflecting the structured causal contribution of this dominant perturbation-aligned axis.

Microbial and metabolite log-ratio balances derived from **V1** exhibited clear separation between diseased and control animals, indicating that **V1** captures a coordinated diet-driven microbial–metabolic shift associated with metabolic dysfunction and HCC progression (Fig. 4D). This consolidation implies that **V1** captures a coordinated diet-driven microbial-metabolic shift resulting in progression toward metabolic dysfunction and, ultimately, HCC. Alignment of **V1** with microbe–metabolite cooccurrence patterns identified by MMvec [36] further linked this factor to bile acid and lipid metabolism, driven in part by *Clostridia*, particularly *Ruminococcus gnavus*. This taxon is a known producer of secondary bile acids via 7*α*-dehydroxylation [37], and its associated metabolites clustered within bile-acid–enriched molecular communities [38] (Fig. 5).

**Fig. 5.**
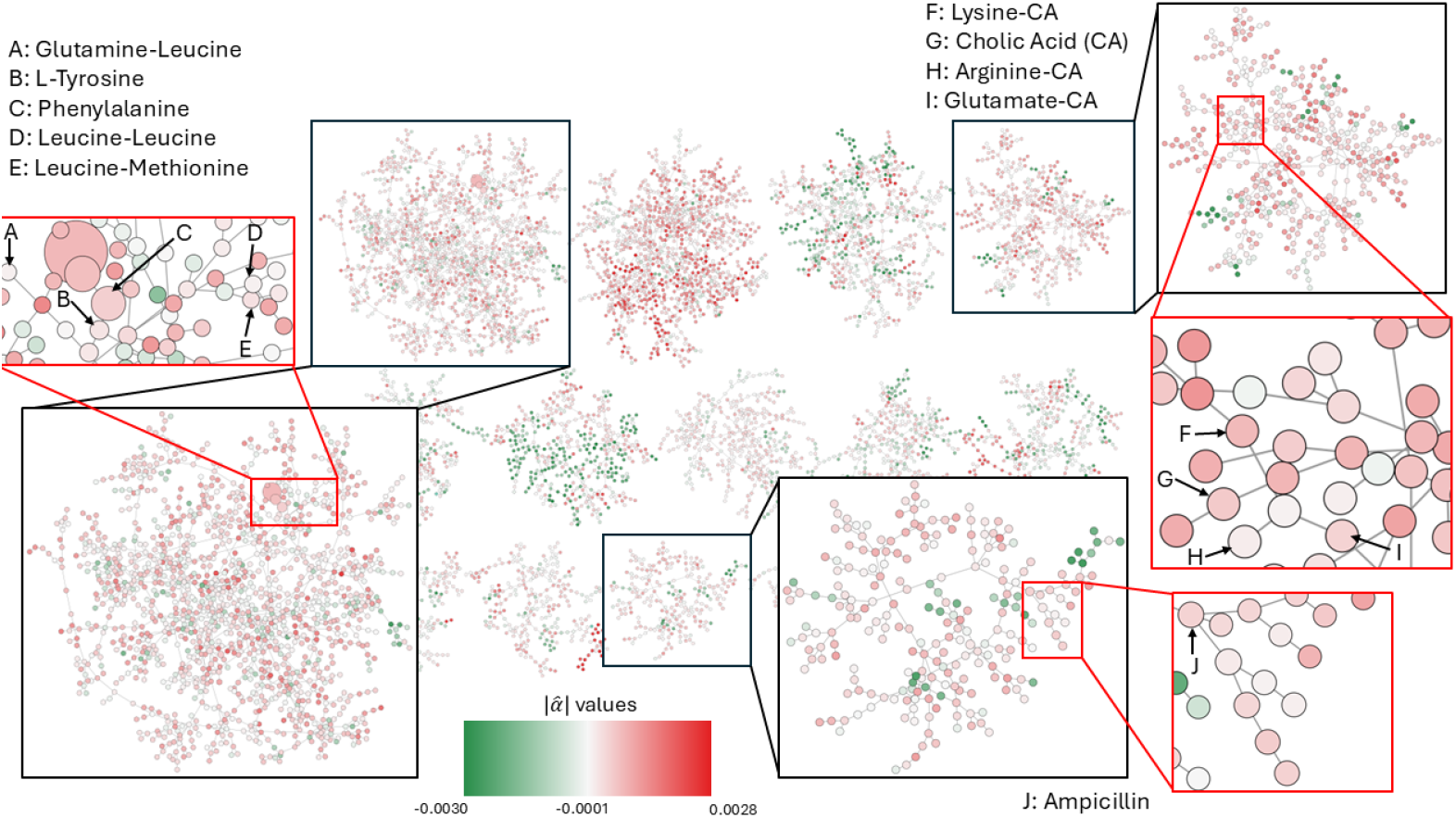
Molecular community network of HCC-associated metabolites. Molecular community network constructed from untargeted metabolomics data. Nodes represent metabolites and edges denote spectral similarity; colors correspond to metabolite’s association with V1, size represents metabolite’s abundance. Examples of metabolites with large absolute causal effects 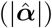 inferred by Factor IV, including bile acids and lipid-related compounds, are highlighted; these molecules occur across specific molecular communities, indicating their structural relatedness.

Although the microbial component highlighted specific taxa, the metabolomic signatures associated with **V1** were distributed across broader molecular families within molecular communities rather than concentrated in individual metabolites [38] (Fig. 5). This pattern indicates that HCC is associated with coordinated shifts in lipid and bile acid metabolism rather than isolated metabolic changes. Together, these results support a mechanistic model in which high-fat diet promotes expansion of *R. gnavus*, driving secondary bile acid production and broader lipid metabolic disruption. Such alterations are consistent with FXR- and TGR5-mediated inflammatory signaling pathways implicated in hepatocarcinogenesis, providing a coherent biological explanation for the strong association between the **V1** factor and HCC-related phenotypes.

To further contextualize these findings, we analyzed the metabolites identified by Factor IV using a molecular community network (MCN) constructed from untargeted metabolomics data. Community-level organization based on spectral similarity revealed that metabolites with strong causal effects ***α*** were distributed across multiple molecular families, including bile acids, lipid derivatives, and antibiotic-associated compounds, rather than appearing as isolated nodes (Fig. 5). In particular, bile acid–enriched communities aligned strongly with ***α*** and separated HCC and obesity phenotypes from controls, consistent with a systems-level perturbation of bile acid metabolism during disease progression [39]. These results reinforce that Factor IV captures biologically structured metabolic variation reflecting coordinated reprogramming of host–microbiome metabolism.

Importantly, this microbe-metabolite axis would have been overlooked using standard differential abundance methods. DESeq2 [23] did not identify *R. gnavus* as significantly associated with HCC, instead highlighting *Enterococcus* species-organisms with high 16S copy number and strong antibiotic resistance [36, 40]. Although these taxa respond robustly to antibiotic treatment, their direct involvement in HCC remains controversial [41]. In contrast, Factor IV isolates the instrument-aligned causal component of microbial variation, allowing HCC-associated microbes and metabolites to be detected even in the presence of strong antibiotic perturbations. This illustrates how IV-based factor modeling can reveal mechanistic pathways that remain inaccessible to association-based analysis.

## 3 Discussion

We present Factor IV, an instrumental variable framework for causal inference in high-dimensional biological systems where endogeneity and unmeasured confounding are intrinsic. In modern multi-omics studies, structured perturbations such as diet, antibiotics, environmental exposures, or genetic variation simultaneously affect large numbers of correlated molecular features, rendering classical IV approaches ill-posed. Factor IV addresses this challenge by learning low-dimensional *instrumental factors* through a sparse low-rank decomposition of the instrument–exposure map, yielding an identifiable first stage even when the number of endogenous variables exceeds the sample size. Under standard exclusion assumptions, these factors remain orthogonal to outcome noise while capturing coordinated biological responses to exogenous perturbations [25, 42].

A central conceptual contribution of Factor IV is the representation of causal effects as a structured decomposition, ***α*** = **V*κ***, where ***κ*** encodes factor-level causal effects and **V** maps these effects onto individual molecular features. This formulation moves beyond feature-wise causal coefficients to model how causal influence propagates through latent biological programs induced by interventions. By aggregating correlated and potentially weak instruments into a small number of stable factors, Factor IV overcomes key limitations of traditional IV estimators, including two-stage least squares and penalized variants, whose performance degrades with many instruments or high-dimensional exposures [43, 44].

Simulation studies confirm that Factor IV accurately recovers both factor-level effects ***κ*** and structured feature-level effects ***α*** under linear and generalized linear models, and remains robust to substantial misspecification, including unobserved interventions and correlated errors. These results establish Factor IV as a statistically principled approach for identifying structured causal effects in complex multi-omic systems.

The biological applications illustrate how these methodological advantages translate to real data. In the HCC mouse model [34], overlapping perturbations—including diet, antibiotics, and immune-related genetic knockouts—jointly influence microbial communities, metabolite profiles, and host phenotypes. Such settings pose a fundamental statistical challenge, as both interventions and responses are high dimensional and strongly correlated. Factor IV resolves this complexity by isolating instrumentaligned latent factors, revealing a dominant perturbation axis driven primarily by diet and antibiotic exposure. The microbial and metabolic loadings on this axis align with bile-acid–mediated inflammatory mechanisms involving taxa such as *Ruminococcus gnavus* and related *Clostridia* [37], patterns obscured by differential abundance analyses that emphasize perturbation responsiveness without causal specificity.

The DIABIMMUNE application highlights a complementary strength of Factor IV in longitudinal, observational settings characterized by rapid ecological change and strong age-dependent confounding [26, 27]. Using age-stratified analyses, Factor IV isolates latent microbial factors whose causal influence is confined to specific developmental windows in infancy. Several of these factors involve taxa previously implicated in immune maturation but masked in association-based analyses once age-driven compositional shifts dominate microbial variation. Aligning microbial structure with exogenous early-life perturbations allows Factor IV to disentangle transient ecological processes from host genetic effects and identify microbial drivers that precede autoantibody seroconversion.

Despite these strengths, Factor IV inherits the core assumptions of instrumental variable analysis, i.e., relevance, exclusion, and independence, which are not empirically testable and require careful scientific justification in each application [42]. Instrument misspecification or weak perturbations may reduce power, and instrumental factors represent aggregated processes rather than single mechanistic pathways, reflecting a trade-off between identifiability and biological resolution. Selection of the latent dimensionality also remains an open challenge in high-dimensional settings.

Looking forward, several extensions are motivated by both statistical and biological considerations, including nonlinear or deep factor models, Bayesian formulations for uncertainty quantification, and incorporation of biological priors such as metabolic or taxonomic structure. More broadly, Factor IV provides a general template for causal inference in emerging multi-modal systems, including longitudinal host–microbiome– metabolite studies and single-cell multi-omics, where high dimensionality, temporal structure, and confounding remain pervasive challenges [20, 35].

## 4 Materials and Methods

### 4.1 Factor Instrumental Variable (Factor IV) Model

We begin with the classical instrumental variables (IV) framework[42, 45], represented conceptually by the directed acyclic graph (DAG) in Fig. 1A. Let **Z** ∈ ℝ^*n×d*^ denote a set of experimental instruments, such as dietary interventions, antibiotic perturbations, or genetic knockouts. Let **X** ∈ ℝ^*n×p*^ represent high-dimensional endogenous variables, for example normalized microbial abundances or metabolite concentrations, and let **Q** ∈ ℝ^*n×t*^ contain observed covariates such as age, weight, or host strain. The outcome of interest is denoted **y** ∈ ℝ^*n*^. At the conceptual data-generating level, we also posit a matrix of unobserved confounders **H** ∈ ℝ^*n×s*^ that simultaneously influence both **X** and **y**. The structural outcome equation takes the form

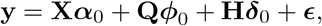

where ***α***_0_ is the causal effect of interest, (***ϕ***_0_, ***δ***_0_) are nuisance coefficients, and ***ϵ*** is a mean-zero structural error. The endogenous regressors arise through the high-dimensional first-stage model

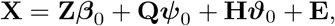

where ***β***_0_ encodes the effects of instruments on **X**, (***ψ***_0_, ***ϑ***_0_) represent observed and unobserved confounding effects, and **E** is a noise matrix.

Endogeneity [42, 46], emerges whenever the endogenous regressors are correlated with the structural error term, that is,

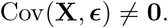

Often, this occurs when the latent confounders **H** influence both **X** and **y**, inducing shared dependence through the terms *f* (**H**) in **X** and *g*(**H**) in **y**. Secondly, correlated noise components in (**E, *ϵ***) naturally arise in biological and technical settings, thereby generating additional dependence. Such phenomena are widespread in multi-omics data; for example, diet variation, inflammation, host immune state, and batch effects all perturb microbial abundance profiles and also influence host outcomes, including metabolic and tumorigenic phenotypes. Because this dependence violates the exogeneity condition required for unbiased ordinary regression, naive regression of **y** on (**X, Q**) is biased and inconsistent [42, 46]. Classical IV estimators such as two-stage least squares[42, 45] typically require more instruments than endogenous regressors and are not directly applicable in modern high-dimensional settings, where both **Z** and **X** may satisfy *d* ≫ *n* and *p* ≫ *n*.

Since **H** is unobserved in practice, IV framework estimation [42] proceeds using the observed-data counterpart of the model. The structural equation is written as

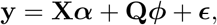

where ***α*** denotes the causal parameter of interest in the observed population. The endogenous regressors satisfy the observable first-stage equation

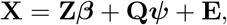

with (***β, ψ***) denoting unknown population parameters. Endogeneity persists through the relation Cov(**X, *ϵ***) = Cov(**E, *ϵ***) ≠ 0, which reflects the latent influence of **H** even though it is not modeled explicitly.

A key challenge in multi-omics studies is that both the instrument matrix **Z** and the endogenous matrix **X** are high-dimensional [14, 25]. Estimating ***β*** ∈ ℝ^*d×p*^ directly is statistically infeasible when *d* ≫ *n*, and the second-stage regression is ill-posed when *p* ≫ *n*. Factor IV addresses this difficulty by imposing a structured low-rank factorization of the instrument-exposure relationship. Specifically, the first-stage coefficient matrix is assumed to satisfy

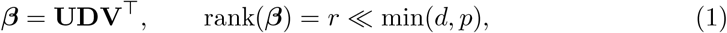

where **U** ∈ ℝ^*d×r*^ and **V** ∈ ℝ^*p×r*^ are sparse factorization matrices and **D** is a diagonal matrix of nonzero singular values. Substituting this decomposition into the first-stage model leads to

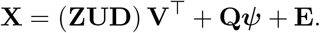

We define the marix **F** = **ZUD** ∈ ℝ^*n×r*^ as the *instrumental factors*: a low-dimensional representation of the exogenous variation in **X** induced by **Z** [25, 47]. Extending the standard IV assumptions, we have

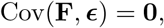

and thus form a valid instrument set for causal identification. A formal proof of the result, i.e., Cov(**F, *ϵ***) = 0, is provided in Supplementary materials Section S3.

Projecting the structural model onto the span of the instrumental factors leads to the reduced second-stage equation

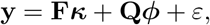

where ***κ*** ∈ ℝ^*r*^ represents the causal effects of the latent factors. The per-feature causal effects are obtained by mapping factor-level effects back into the original feature space according to

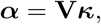

so that each microbial or metabolite feature inherits its causal effect from its loading in **V**. See Supplementary Material Corollary S4 for the formal validity and identifiability of the per-feature causal parameters ***α*** = **V*κ***.

Estimation proceeds by fitting a sparse low-rank regression of **X** on (**Z, Q**) using Gaussian co-sparse factor models appropriate for continuous data[19, 20, 35]. These methods yield estimates 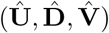 and therefore estimated instrumental factors 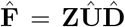. A second-stage IV or GMM regression of **y** on 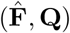 provides an estimate of ***κ***, and consequently the per-feature causal effects are given by 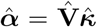. Factor IV framework provides a principled and interpretable way to (i) extract latent perturbation-driven variation from high-dimensional multi-omics data[48–50], (ii) construct valid instruments in the presence of unobserved confounding, and (iii) estimate causal effects of microbial and metabolic features on host phenotypes even when the number of endogenous variables far exceeds the sample size.

#### Generalized linear first-stage model for high-dimensional count data

In many multi-omics applications, including microbiome studies, the endogenous variables **X** are not continuous but binary/count-valued (e.g., 16S rRNA read counts, metagenomic counts, or RNA-seq-type abundances). In such settings, a Gaussian working model for the first stage is inappropriate, and it is more natural to work within a generalized linear model (GLM) framework with Bernaulli or Poisson or negative binomial (NB) likelihoods. The Factor IV framework extends directly to this setting by combining high-dimensional IV structure with low-rank generalized co-sparse factor regression [20, 35].

For sample *i* ∈ {1, …, *n*} and feature (taxon/metabolite) *j* ∈ {1, …, *p*}, we model

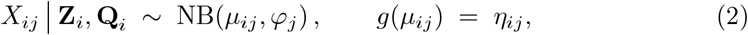

where *µ*_*ij*_ = 𝔼 [*X*_*ij*_ | **Z**_*i*_, **Q**_*i*_] is the conditional mean, *φ*_*j*_ *>* 0 is a feature-specific dispersion parameter, and *g*(·) is the log link, *g*(*µ*) = log *µ*. In the NB parameterization we use,

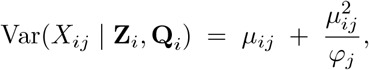

which allows overdispersion relative to the Poisson model, as is typical for microbiome and other sequencing data. A Poisson first-stage model is obtained as a special case by letting *φ*_*j*_ → ∞. We define the linear predictor linked to the mean parameter ***µ*** = [*µ*_*ij*_] as

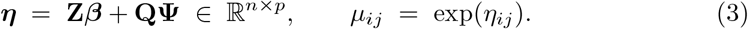

To encode interpretable low-dimensional structure in the instrument-exposure map in the high-dimensional regime, we impose a sparse low-rank factorization on the coefficient matrix ***β*** (1) and rewrite (3) as

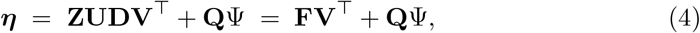

where the *r*-dimensional *instrumental factors* **F** = **ZUD** summarize the low-dimensional, instrument-induced component of the high-dimensional **X**.

In this GLM first-stage model, conditional on (**Z, Q**), the distribution of **X** depends on the instruments only through the low-dimensional factors **F**. Under the standard IV assumptions imposed at the level of the observed instruments **Z**- (i) *relevance*: **Z** is associated with **X** through ***β***; (ii) *exclusion*: **Z** affects **y** only through **X** (given **Q**); and (iii) *independence*: **Z** ⊥ ***ϵ*** | **Q**. This implies that the factors inherit exogeneity:

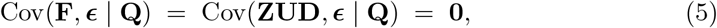

so that **F** (together with **Q**) can be used as a valid low-dimensional instrument set in the second-stage outcome model. Please find a formal proof of the exogeneity of the instrumental factors in Lemma 1 of the Supplementary Materials.

The second-stage model may itself be specified as a linear or generalized linear regression. For a continuous outcome we take

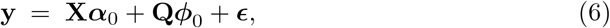

with endogeneity arising from Cov(**X, *ϵ***) ≠ 0. In a GLM formulation (e.g., for binary or count outcomes) we instead write

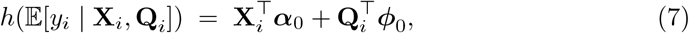

with canonical link *h*( *·*) (e.g., logit or log), and then replace **X** by its instrumented low-rank component encoded by **F**. In practice, we first estimate (**U, D, V**) by fitting a generalized co-sparse factor regression of **X** on (**Z, Q**) using the `gofar` or `nbfar` implementations for Poisson or NB responses [20, 35]. This yields estimated factors 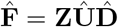, on which we perform a second-stage IV or GMM regression

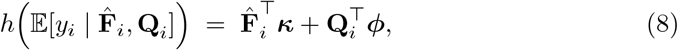

to obtain factor-level effects 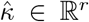. Finally, per-feature causal effects are reconstructed via

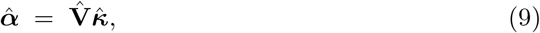

providing a high-dimensional estimate of the causal influence of individual microbial or metabolite features on the outcome while leveraging a low-dimensional, instrumentaligned representation of the exposure.

#### Multi-omics integration and identifiability

In many biological systems, instruments may influence outcomes through multiple molecular layers that are only partially observed in single-omics analyses. Expanding the endogenous feature space to include additional omics modalities can therefore strengthen the plausibility of the exclusion restriction by capturing mediating pathways that would otherwise remain unmeasured. In the Factor IV framework, multi-omics measurements improve causal identification by allowing instrument-aligned latent factors to absorb a larger fraction of exogenous biological variation. A formal justification of this effect, using a causal graph-based argument, is provided in SupplementaryMaterials Section S6.

### 4.2 Simulation study design

We conducted a set of simulation experiments to (i) verify that Factor IV recovers the ground-truth causal effects ***α***_0_ when data are generated from the proposed model, and (ii) assess robustness to endogeneity arising from unobserved confounding or correlated errors in high-dimensional settings for both continuous and count-valued endogenous variables.

#### Generative setup and latent structure

All simulations were generated from the observed-data Factor IV model described in Section 4.1. We considered *n* samples, *d* instruments, *p* endogenous variables, *t* observed confounders, and *k* latent instrumental factors, with *p* ≫ *n* and *k* ≪ min {*d, p*}. In the experiments underlying Fig. 2, we set (*n, d, p, t, k*) = (200, 20, 100, 3, 3) and used the diagonal factor-strength matrix **D** = diag(3, 4, 5). Instruments and confounders were generated independently as *Z*_*ij*_ ~ 𝒩 (0, 1), and *Q*_*ij*_ ~ 𝒩 (0, 1).

The low-rank instrument-exposure map was defined as ***β*** = **UDV**^⊤^ ∈ ℝ^*d×p*^, where **U** ∈ ℝ^*d×k*^ and **V** ∈ ℝ^*p×k*^ are sparse [19, 20]. Specifically, **U** and **V** were generated with entries from Unif(−1, 1) and sparsified by hard thresholding: *U*_*ℓr*_ ← 0 if |*U*_*ℓr*_| *<* 0.75, *V*_*jr*_ ← 0 if |*V*_*jr*_| *<* 0.75. Columns were then renormalized to satisfy **V**^⊤^**V** = **I**_*k*_. To satisfy the condition **U**^⊤^**X**^⊤^**XU***/n* = **I**_*k*_ in the continuous-Gaussian setting, we generated **X** conditional on **U** using the orthogonal completion and covariance projection algorithm implemented in the function `simulate_X_given_U()`, which constructs **X** so that the sample covariance of **X** in the directions of **U** matches the identity [19, 20].

#### Endogeneity mechanisms

We introduced endogeneity via unobserved confounding or correlated errors, each controlled by *λ* ∈ [0, 1]. These mechanisms are formally equivalent under the observed-data model and induce identical violations of exogeneity (Supplementary Materials Section S8).

##### a) Latent confounder mechanism

A latent scalar confounder **H** ~ 𝒩_*n*_(0, **I**_*n*_) entered both the first- and second-stage signals:

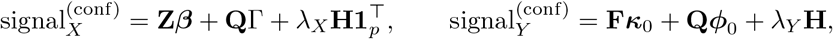

with 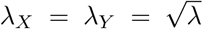. Because **H** is unobserved, this induces Cov(**X, *ϵ***) ≠ 0, violating the exclusion restriction.

##### b) Correlated-error mechanism

We generated noise components *e*_1_, *e*_2_ ~ 𝒩_*n*_(0, **I**_*n*_) and **E**_3_ ~ 𝒩_*n×p*_(0, 1), and constructed raw errors

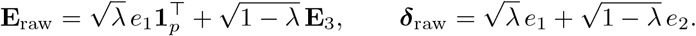

When *λ >* 0, **X** and ***ϵ*** share the common component *e*_1_, again giving Cov(**X, *ϵ***) ≠ 0.

#### First-stage models and signal-to-noise control

We generated the observed exposures by combining the structured first-stage signal with noise scaled to achieve a target signal-to-noise ratio. In the continuous (Gaussian) setting, we first drew a confounder-loading matrix Γ ~ 𝒩_*t×p*_(0, 1) and constructed the baseline signal signal_*X*_ = **Z*β*** + **Q**Γ, or its confounder-modified version when a latent endogeneity mechanism was active. Raw noise **E**_raw_ was then scaled according to 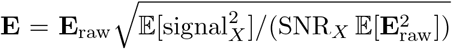 with SNR_*X*_ = 3, ensuring that Var(signal_*X*_)*/*Var(**E**) ≈ SNR_*X*_. The observed Gaussian exposure matrix was finally set to **X** = signal_*X*_ + **E**.

In the negative binomial first stage, the same linear predictor served as the NB mean structure, now written as *η*_*ij*_ = (signal_*X*_)_*ij*_ with *µ*_*ij*_ = exp(*η*_*ij*_), followed by draws *X*_*ij*_ | **Z**_*i*_, **Q**_*i*_ ~ NB(*µ*_*ij*_, *φ*) using a fixed dispersion parameter *φ* = 2. This specification preserved the intended low-rank latent factor geometry while incorporating realistic overdispersion characteristic of microbiome sequencing counts. The log link ensured that instrument-induced perturbations in **Z** propagated multiplicatively into the conditional mean structure, mirroring how sequencing read counts respond to biological interventions.

Outcome generation followed a parallel design. Factor-level causal effects were drawn as ***κ***_0_ ~ Unif(2, 4)^*k*^ and mapped to per-feature causal effects via ***α***_0_ = **V*κ***_0_, with confounder effects sampled as ***ϕ***_0_ ~ 𝒩_*t*_(0, **I**_*t*_). The structured outcome signal was defined as signal_*Y*_ = **F*κ***_0_ + **Q*ϕ***_0_, or its endogeneity-augmented version when hidden confounders were present. Noise was scaled analogously using 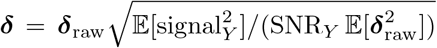with SNR_*Y*_ = 3, and the final outcome was computed as **y** = signal_*Y*_ + ***δ***. This design preserved the latent factor structure while controlling the difficulty of causal effect recovery across simulation scenarios.

#### Estimation and benchmarking

Factor IV was implemented exactly as in the observed-data framework. For each simulated dataset, the first stage was estimated using GOFAR for Gaussian exposures [20] or NBFAR for negative binomial exposures [35], producing estimates 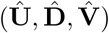 and corresponding instrumental factors defined as 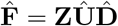. The second stage regressed the outcome on these estimated factors and observed confounders using the model 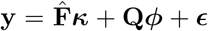, from which factor-level coefficients 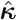were obtained and mapped into per-feature causal estimates using 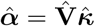. This two-step procedure ensured that only the variation in **X** driven by the instruments was used to estimate causal effects, consistent with the standard IV logic but adapted to the low-rank, high-dimensional setting.

Benchmarking included methods that ignore the instrumental variable structure. For continuous exposures, we applied feature-wise naive linear regression, regressing the outcome on each column of **X** and **Q** and taking the coefficient of the exposure as the estimate of its causal effect. For count-valued exposures, we used DESeq2 [23], treating the outcome as a covariate in the feature-wise negative binomial GLM and extracting the coefficient associated with the phenotype as a pseudo-causal effect. These baselines illustrate the bias that arises when endogeneity is ignored.

#### Evaluation metrics

Performance was assessed across Monte Carlo replicates using several complementary metrics. Accuracy of causal effect recovery was measured using the mean squared error 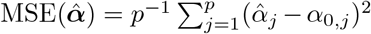 and the coefficient of determination 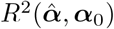 summarizing correlation between estimated and true effects. Prediction accuracy was evaluated through 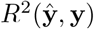 and through the ability to reconstruct the latent perturbation signal using 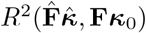. When the data-generating process induced sparse causal effects, we additionally computed sensitivity and specificity for correctly identifying nonzero entries of ***α***_0_ under multiple-testing correction. All simulation results are presented in Fig. 2.

### Application to a Hepatocellular Carcinoma Mouse Model

#### Dataset description

We analyzed a multi-omics dataset from a murine hepatocellular carcinoma (HCC) study reported by Shalapour et al. [34]. The dataset integrates gut microbiome profiles, untargeted metabolomics, immune phenotypes, dietary interventions, antibiotic treatments, and multiple genetic knockouts designed to perturb host-microbiome interactions during tumor development. Each mouse was profiled across microbial, metabolic, and phenotypic modalities, yielding a high-dimensional endogenous feature space with multiple experimentally controlled perturbations.

#### Data preprocessing

Microbiome data were processed at the ASV level. Raw count tables were filtered to remove rare taxa and normalized using centered log-ratio (CLR) transformation after adding a small pseudocount, yielding approximately Gaussian features suitable for linear modeling [51]. Metabolomics data were log-transformed and normalized by total ion count to account for global intensity differences across samples. The resulting microbiome and metabolite features were concatenated to form a single endogenous matrix **X** ∈ ℝ^*n×p*^. Host phenotypes, including HCC status, obesity, non-alcoholic steatohepatitis (NASH), IgA abundance, and B-cell depletion, were encoded as binary or continuous outcomes depending on the analysis.

#### Instrumental variables and observed confounders

Instrumental variables **Z** ∈ ℝ^*n×d*^ were defined using experimentally assigned perturbations: dietary intervention (high-fat diet versus control), antibiotic treatments (vancomycin, neomycin, ampicillin, and metronidazole), and immune-related genetic knockouts (including IgA^−*/*−^, IgAkMUP, MupHe, B6, and STAM backgrounds). These perturbations directly affect microbial and metabolic composition and were treated as exogenous with respect to host disease outcomes conditional on the endogenous variables. Observed confounders **Q** ∈ ℝ^*n×t*^ included age, body weight, and mouse strain, which were included in all stages of the analysis.

#### Factor IV model

We modeled the high-dimensional endogenous feature matrix using the observed-data Factor IV first-stage specification **X** = **ZUDV**^⊤^ + **Q*ψ*** + **E**, where **U** ∈ ℝ^*d×r*^ and **V** ∈ ℝ^*p×r*^ are sparse loading matrices, **D** is a diagonal matrix of singular values, and *r* ≪ min(*d, p*). The model was fit using generalized co-sparse factor regression with a Gaussian working likelihood, as implemented in the `gofar` package [20]. The rank was fixed at *r* = 2 based on explained variance and stability across bootstrap replicates. Estimated instrumental factors were constructed as 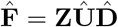.

For each host phenotype *y*, we estimated causal effects by fitting the second-stage regression model 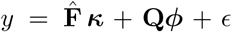, where ***κ*** captures factor-level causal effects and ***ϕ*** adjusts for observed confounders. For continuous outcomes, estimation was performed using ordinary least squares; for binary outcomes, a logistic regression model was used. Feature-level causal effects were reconstructed as 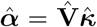, yielding a per-feature estimate aligned with the instrument-induced latent structure.

#### Comparison with association-based methods

For benchmarking, we applied differential abundance analysis using DESeq2 [23] to the microbiome count data. Feature-wise negative binomial models were fit with the host phenotype as a covariate, adjusting for sequencing depth and multiple testing. DESeq2 results were used solely as an association-based reference and were not interpreted causally.

#### Statistical inference and implementation

All analyses were conducted in R. Factor IV model fitting used the `gofar` [20] package for first-stage estimation and standard regression routines for second-stage inference. Statistical significance of reconstructed feature-level effects was assessed using Waldtype statistics derived from the second-stage models. All code used for preprocessing, model fitting, and analysis is provided in the accompanying reproducibility repository.

##### Generation of a molecular community network

An unpruned molecular network was generated in which all pairwise node connections were retained. Network construction parameters were specified using the Advanced Network Options in the GNPS environment [52] as follows: minimum pairs cosine similarity of 0.1, Network TopK set to 100, and maximum connected component size set to 0.

The resulting unpruned network served as the input for Molecular Community Network (MCN) analysis. The unpruned .`graphml` file was downloaded from GNPS and processed using the MCN workflow to generate a corresponding .`graphml` file representing the derived community network. Community detection was performed using the Louvain community detection algorithm, which partitions the network by maximizing modularity. Following community assignment, intra-community connections were pruned using a Maximum Weight Spanning Tree (MWST) algorithm, retaining only the highest-weight edges while preserving a single connected path within each community [38]. The complete GNPS workflow is publicly available at https://gnps.ucsd.edu/ProteoSAFe/status.jsp?task=7f37d16b6d034aed8219fd8f6fad8a8e.

### Longitudinal Infant Microbiome Analysis in the DIABIMMUNE Cohort

We applied Factor IV to the longitudinal DIABIMMUNE infant cohort [26, 27], which studies how early-life gut microbiome development relates to genetic risk for type 1 diabetes (T1D) and the emergence of islet autoantibodies. The cohort follows infants from birth through early childhood and includes dense microbiome sampling together with host genetics, environmental exposures, and immunological outcomes. Compared to the HCC application, this dataset introduces additional challenges, including repeated measurements, strong age dependence, compositional microbiome data, and high-dimensional sparsity.

#### Data Processing

Microbiome data were generated using both 16S rRNA (V4) and metagenomic sequencing, producing ASV-level and species-level count tables, respectively. Host metadata included HLA DR3/DR4 risk haplotypes, delivery mode, feeding mode, early-life environmental exposures (e.g., siblings and day-care), and multiple autoimmune outcomes, including seroconversion to islet autoantibodies and T1D diagnosis. Standard quality control and prevalence filtering were applied to microbiome count tables. To mitigate compositional effects, microbial features were analyzed on a normalized scale, and participant-level covariates were retained as observed confounders.

To accommodate the strong age dependence of infant gut microbiome development, we performed age-stratified Factor IV analyses instead of fitting a single longitudinal model. This reduces confounding from age-driven compositional shifts and avoids within-subject dependence. Repeated-measure samples were partitioned into three developmental windows: early (0–12 months), mid (12–24 months), and late (*>* 24 months). Within each window, we constructed a cross-sectional dataset by retaining, for each participant, the most recent sample in that interval, yielding three observed datasets 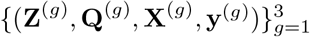 with at most one observation per participant per group.

For each age-stratified analysis, microbiome count tables were merged with participant-level metadata and autoimmune outcomes using participant identifiers. Modality-specific depth filtering was applied separately to metagenomic and 16S data, features with zero prevalence were removed, and the resulting participant-by-feature matrices **X**^(*g*)^ were paired with outcome vectors **y**^(*g*)^ and observed covariates. The instrument matrix **Z**^(*g*)^ was constructed from early-life variables that plausibly perturb microbiome assembly while having no direct effect on autoimmune outcomes except through microbial pathways, including host HLA risk category, delivery mode (vaginal versus Cesarean), infant feeding mode (breastfeeding versus formula), and household environmental exposures such as siblings, pets, and day-care attendance. These variables are well-established drivers of early microbiome colonization and are commonly treated as exogenous with respect to downstream autoimmune phenotypes conditional on microbial composition [26, 27, 42]. Observed confounders **Q**^(*g*)^ captured demographic, anthropometric, and technical factors that may influence both microbiome composition and autoimmune risk, including infant sex, country or study site, sequencing batch, growth-related measures (height-for-age and BMI-for-age *z*-scores), and age (in months) at sampling. All instruments and confounders were standardized and included explicitly in both stages of the Factor IV model, ensuring that causal identification was driven by exogenous perturbation-aligned variation rather than demographic or technical effects.

#### Factor IV Analysis

For each developmental window, we applied age-stratified Factor IV using a generalized co-sparse factor regression framework. Within each age group *g*, the endogenous microbiome matrix **X**^(*g*)^ (CLR-transformed for metagenomics and 16S data) was regressed on the combined design matrix [**Q**^(*g*)^, **Z**^(*g*)^], where observed confounders and instruments were jointly included but indexed separately for causal identification. First-stage factor estimation was performed using GOFAR [20], with a Gaussian working likelihood for CLR-transformed features and a fixed candidate rank of *r* = 5, chosen to capture dominant perturbation-driven structure while remaining conservative relative to sample size. Strong sparsity was enforced on both instrument loadings and feature loadings (with sparsity levels `spU` = 0.99 and `spV` = 0.98) to promote interpretability and stability in the high-dimensional setting. Model fitting used a grid of 30 regularization parameters with five-fold cross-validation, and convergence tolerances were tightened (maximum iterations = 5000-10000) to ensure numerical stability. Random seeds were fixed across age groups for reproducibility. Estimated instrumental factors were constructed as 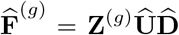 and carried forward to the second stage, where causal effects on autoimmune phenotypes were estimated via generalized linear regression of the form 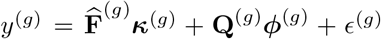. Feature-level causal effects were reconstructed as 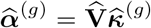, enabling age-specific attribution of microbial taxa to downstream immune outcomes.

## Data Availability

All data used in this study are publicly available. The hepatocellular carcinoma (HCC) mouse multi-omics dataset analyzed here was obtained from the original published study [34]. The DIABIMMUNE longitudinal infant microbiome data were downloaded from MicrobiomeDB (https://microbiomedb.org) [26, 27]. Processed data objects, analysis-ready feature tables, and selected intermediate outputs from the Factor IV analyses for both the HCC and DIABIMMUNE applications are available via a Zenodo repository at https://doi.org/10.5281/zenodo.17968386. For the HCC metabolomics analysis, molecular networking and Molecular Community Network (MCN) intermediate outputs were generated using the GNPS platform and are publicly accessible through the workflow link https://gnps.ucsd.edu/ProteoSAFe/status.jsp?task=7f37d16b6d034aed8219fd8f6fad8a8e [38].

## Code Availability

All code used for simulations, data preprocessing, model fitting, and downstream analyses is publicly available at https://github.com/amishra-stats/FactorIV.

The repository provides a modular and reproducible implementation of the Factor IV framework, including (i) simulation studies, (ii) analysis of the DIABIMMUNE longitudinal cohort, and (iii) analysis of the hepatocellular carcinoma (HCC) mouse model. Each component contains documented R scripts for data preparation, factor estimation, causal analysis, and visualization, together with example outputs and figure-generation code.

The analyses rely on the `gofar` and `nbfar` R packages for fitting sparse low-rank Gaussian and negative binomial factor regression models, respectively. Both packages are open source and available via GitHub, with installation instructions provided in the Factor IV repository. All software versions and package dependencies required to reproduce the results are specified in the repository documentation.

## Code Availability

All code for simulations, data preprocessing, model fitting, and downstream analyses is available at https://github.com/amishra-stats/FactorIV. The repository contains fully reproducible workflows for the simulation studies, the DIABIMMUNE longitudinal analysis, and the hepatocellular carcinoma (HCC) application, including scripts for data preparation, factor estimation, causal modeling, and figure generation. Sparse low-rank first-stage models are implemented using the `gofar` and `nbfar` R packages, available at https://github.com/amishra-stats/gofar and https://github.com/amishra-stats/nbfar/tree/master, respectively. Installation instructions and software dependencies are provided in the repository.

## Author Contributions

A.M. and J.M. created the idea for the work and conducted all the data analysis. A.A. conducted metabolomics interpretation L.C. constructed molecular networks for the HCC analysis. A.M., J.M., and A.A. wrote the manuscript. J.M., L.C., A.A., and R.B. reviewed and edited the manuscript. R.B. provided senior oversight of the project.

## Acknowledgements

We thank Dean Eckles for helpful discussions and feedback on the manuscript.

