## Supplemental Materials for "Moving from Association to Causation: Instrumental factor models for causal inference in high-dimensional multi-omics data"

#### S1. Observed-Data Model and IV Assumptions

We work throughout with the observed-data formulation used in the main manuscript. Let

$$\mathbf{y} \in \mathbb{R}^n, \quad \mathbf{X} \in \mathbb{R}^{n\times p}, \quad \mathbf{Z} \in \mathbb{R}^{n\times d}, \quad \mathbf{Q} \in \mathbb{R}^{n\times t}$$

denote, respectively, the outcome vector, high-dimensional endogenous regressors, instruments, and observed confounders.

The structural outcome model is

$$\mathbf{y} \;=\; \mathbf{X}\boldsymbol{\alpha}_0 \;+\; \mathbf{Q}\boldsymbol{\phi}_0 \;+\; \boldsymbol{\epsilon}, \tag{1}$$

where $\boldsymbol{\alpha}_0 \in \mathbb{R}^p$ is the causal parameter of interest, $\boldsymbol{\phi}_0 \in \mathbb{R}^t$ captures the effect of observed confounders, and $\boldsymbol{\epsilon} \in \mathbb{R}^n$ is a mean-zero structural error, $\mathbb{E}[\boldsymbol{\epsilon}] = \mathbf{0}$.

The endogenous regressors $\mathbf{X}$ are generated from a high-dimensional first-stage model

$$\mathbf{X} \;=\; \mathbf{Z}\boldsymbol{\beta} \;+\; \mathbf{Q}\boldsymbol{\psi} \;+\; \mathbf{E}, \tag{2}$$

where $\boldsymbol{\beta} \in \mathbb{R}^{d\times p}$ is the instrument–exposure coefficient matrix, $\boldsymbol{\psi} \in \mathbb{R}^{t\times p}$ encodes the effect of observed confounders on $\mathbf{X}$, and $\mathbf{E} \in \mathbb{R}^{n\times p}$ is a noise matrix.

Endogeneity arises whenever

$$\mathrm{Cov}(\mathbf{X}, \boldsymbol{\epsilon}) \;\neq\; \mathbf{0}, \tag{3}$$

which we interpret as $\mathrm{Cov}(\mathbf{E}, \boldsymbol{\epsilon}) \neq 0$. In multi-omics applications, this typically reflects unobserved biological or technical sources of variation—for example, diet, inflammation, or batch effects—that simultaneously perturb both $\mathbf{X}$ and $\mathbf{y}$. Under such conditions, a naive regression of $\mathbf{y}$ on $(\mathbf{X}, \mathbf{Q})$ yields biased and inconsistent estimates of $\boldsymbol{\alpha}_0$ [2, 5, 3].

The classical IV framework [1, 3, 6] imposes the following assumptions at the level of the observed instruments $\mathbf{Z}$:

*Department of Statistics, University of Georgia, Athens, GA, USA..
†Department of Biology, New York University, New York, NY, USA.
‡Department of Pharmaceutical Sciences, University of Connecticut, Storrs, CT, USA.
§Department of Statistics, University of Georgia, Athens, GA, USA..
¶Center for Genomics and Systems Biology, New York University; Prescient Design, Genentech. Co-senior author.
‖Gutz Analytics, Boulder, CO, USA. Contributed equally.

**(IV1) Relevance.** Instruments must explain nontrivial variation in the endogenous regressors. Formally, there exists a nonzero matrix $\boldsymbol{\beta}$ such that

$$\operatorname{rank}\big(\mathbb{E}[\mathbf{Z}^\top \mathbf{X}]\big) \;=\; r > 0.$$

In the Factor IV setting, this translates to $\operatorname{rank}(\mathbf{Z}\boldsymbol{\beta}) = r$, where $r$ is the factor rank.

**(IV2) Exclusion Restriction.** Instruments affect the outcome only through the endogenous regressors and observed confounders; they have no direct effect on $\mathbf{y}$. In the structural model (1), this is encoded by the absence of $\mathbf{Z}$ among the regressors and by the orthogonality condition

$$\mathbb{E}[\mathbf{Z}^\top \boldsymbol{\epsilon}] = \mathbf{0}.$$

**(IV3) Independence (or Instrument Exogeneity).** Instruments are independent of the structural error and the noise in the endogenous regressors:

$$\mathbb{E}[\mathbf{Z}^\top \boldsymbol{\epsilon}] = \mathbf{0}, \qquad \mathbb{E}[\mathbf{Z}^\top \mathbf{E}] = \mathbf{0}.$$

Equivalently, any unobserved confounders may influence $(\mathbf{X}, \mathbf{y})$ but not $\mathbf{Z}$, a standard assumption in the IV literature [3].

**(IV4) Rank Condition (Identification).** The matrix of IV moments must be of full column rank:

$$\operatorname{rank}\Big(\mathbb{E}\Big[(\mathbf{Z}, \mathbf{Q})^\top \mathbf{X}\Big]\Big) \;=\; p,$$

or in the reduced Factor IV representation, the analogous rank condition for the factor regressors and instruments. This guarantees identification of $\boldsymbol{\alpha}_0$ via generalized method of moments (GMM) or two-stage least squares (2SLS) [1, 6].

We next show how these classical assumptions extend to the Factor IV setting and validate the use of *instrumental factors* as low-dimensional instruments.

### S2. Instrumental Factors in Factor IV

Factor IV imposes a sparse low-rank structure on the high-dimensional coefficient matrix $\boldsymbol{\beta}$:

$$\boldsymbol{\beta} \;=\; \mathbf{U}\mathbf{D}\mathbf{V}^\top, \qquad \operatorname{rank}(\boldsymbol{\beta}) = r \ll \min\{d, p\}, \tag{4}$$

where $\mathbf{U} \in \mathbb{R}^{d \times r}$ and $\mathbf{V} \in \mathbb{R}^{p \times r}$ are sparse loading matrices and $\mathbf{D} = \operatorname{diag}(d_1, \ldots, d_r)$ contains nonzero singular values.

We define the *instrumental factors* as

$$\mathbf{F} \;=\; \mathbf{Z}\mathbf{U}\mathbf{D} \;\in\; \mathbb{R}^{n \times r}, \tag{5}$$

which represent the low-dimensional variation in $\mathbf{X}$ that is driven by $\mathbf{Z}$. Substituting (4) into the first-stage model (2) yields

$$\mathbf{X} \;=\; \mathbf{Z}\mathbf{U}\mathbf{D}\mathbf{V}^\top \;+\; \mathbf{Q}\boldsymbol{\psi} \;+\; \mathbf{E} \;=\; \mathbf{F}\mathbf{V}^\top \;+\; \mathbf{Q}\boldsymbol{\psi} \;+\; \mathbf{E}. \tag{6}$$

Thus, the high-dimensional instrument–exposure relationship is mediated through $r$ latent factors $\mathbf{F}$.

The main manuscript states that, under (IV1)–(IV3), the instrumental factors are valid instruments in the sense that

$$\mathrm{Cov}(\mathbf{F}, \boldsymbol{\epsilon}) = \mathbf{0}.$$

We formalize this below.

### S3. Validity of Instrumental Factors

**Theorem 1** (Instrumental Validity of the Factor IV Regressors)**.** *Suppose the observed-data model* (1)–(2) *holds and the IV assumptions (IV1)–(IV3) are satisfied. Let* $\boldsymbol{\beta} = \boldsymbol{U}\boldsymbol{D}\boldsymbol{V}^\top$ *be the sparse low-rank factorization* (4)*, and define the instrumental factors* $\boldsymbol{F} = \boldsymbol{Z}\boldsymbol{U}\boldsymbol{D}$ *as in* (5)*. Then*

$$\mathrm{Cov}(\boldsymbol{F}, \boldsymbol{\epsilon}) = \mathbf{0},$$

*so that* $\boldsymbol{F}$ *(together with* $\boldsymbol{Q}$*) forms a valid instrument set for the structural model* (1)*.*

*Proof.* By definition of $\mathbf{F}$,

$$\mathrm{Cov}(\mathbf{F}, \boldsymbol{\epsilon}) = \mathrm{Cov}(\mathbf{ZUD}, \boldsymbol{\epsilon}).$$

Using the linearity of covariance in its first argument and the fact that $(\mathbf{U}, \mathbf{D})$ are deterministic matrices, we obtain

$$\mathrm{Cov}(\mathbf{F}, \boldsymbol{\epsilon}) = \mathbf{UD}\,\mathrm{Cov}(\mathbf{Z}, \boldsymbol{\epsilon}).$$

Under the exclusion and independence assumptions (IV2)–(IV3), we have

$$\mathbb{E}[\mathbf{Z}^\top \boldsymbol{\epsilon}] = \mathbf{0} \quad \Longrightarrow \quad \mathrm{Cov}(\mathbf{Z}, \boldsymbol{\epsilon}) = \mathbf{0},$$

because $\mathbb{E}[\boldsymbol{\epsilon}] = \mathbf{0}$ by construction. It follows that

$$\mathrm{Cov}(\mathbf{F}, \boldsymbol{\epsilon}) = \mathbf{UD} \cdot \mathbf{0} = \mathbf{0}.$$

Thus, no variation in $\mathbf{F}$ is explained by the structural error $\boldsymbol{\epsilon}$, and the instrumental factors inherit the exogeneity and exclusion properties of $\mathbf{Z}$. Relevance follows from (IV1) and the fact that $\mathrm{rank}(\mathbf{Z}\boldsymbol{\beta}) = \mathrm{rank}(\mathbf{F}\mathbf{V}^\top) = r > 0$. Hence, $\mathbf{F}$ (along with $\mathbf{Q}$) can serve as a valid instrument set in the second-stage IV regression for $\boldsymbol{\alpha}_0$ [3, 6]. □

### S4. Per-Feature Causal Effects and Factor-Level Representation

In the Factor IV framework, causal effects are first estimated at the level of latent factors and then mapped back to per-feature effects. We now formalize this and give a detailed proof that the resulting per-feature parameter vector satisfies the IV moment conditions.

#### S4.1 Factor-Level Structural Equation

Starting from the first-stage factor representation (6),

$$\mathbf{X} = \mathbf{F}\mathbf{V}^\top + \mathbf{Q}\boldsymbol{\psi} + \mathbf{E},$$

we substitute this into the structural outcome model (1):

$$\begin{aligned}\mathbf{y} &= \big(\mathbf{F}\mathbf{V}^\top + \mathbf{Q}\boldsymbol{\psi} + \mathbf{E}\big)\boldsymbol{\alpha}_0 + \mathbf{Q}\boldsymbol{\phi}_0 + \boldsymbol{\epsilon} \\ &= \mathbf{F}\underbrace{\mathbf{V}^\top\boldsymbol{\alpha}_0}_{\boldsymbol{\kappa}_0} + \mathbf{Q}(\boldsymbol{\psi}\boldsymbol{\alpha}_0 + \boldsymbol{\phi}_0) + \underbrace{\mathbf{E}\boldsymbol{\alpha}_0 + \boldsymbol{\epsilon}}_{\mathbf{u}},\end{aligned}$$

where we define the factor-level causal parameter

$$\boldsymbol{\kappa}_0 := \mathbf{V}^\top\boldsymbol{\alpha}_0 \in \mathbb{R}^r$$

and the composite error term

$$\mathbf{u} := \mathbf{E}\boldsymbol{\alpha}_0 + \boldsymbol{\epsilon}.$$

Let

$$\boldsymbol{\gamma}_0 := \boldsymbol{\psi}\boldsymbol{\alpha}_0 + \boldsymbol{\phi}_0 \in \mathbb{R}^t.$$

Then the outcome admits the reduced representation

$$\mathbf{y} = \mathbf{F}\boldsymbol{\kappa}_0 + \mathbf{Q}\boldsymbol{\gamma}_0 + \mathbf{u}. \tag{7}$$

This is a standard linear IV structure with regressors $(\mathbf{F}, \mathbf{Q})$, factor-level causal parameters $(\boldsymbol{\kappa}_0, \boldsymbol{\gamma}_0)$, and error $\mathbf{u}$.

#### S4.2 Moment Conditions and Identification

We now show that the per-feature causal parameter $\boldsymbol{\alpha}_0$ obtained by mapping back from factor space satisfies the IV orthogonality conditions.

**Corollary 1** (Validity and Identification of Per-Feature Causal Effects)**.** *Assume the observed-data model* (1)–(2)*, the factor decomposition* (4)*, and the IV conditions (IV1)–(IV4). Let $\boldsymbol{F}$ be the instrumental factors defined in* (5)*, and let the factor-level structural model be given by* (7)*. Suppose additionally that*

$$\mathbb{E}[\boldsymbol{F}^\top\mathbf{u}] = \mathbf{0}, \qquad \mathbb{E}[\boldsymbol{Q}^\top\mathbf{u}] = \mathbf{0},$$

*i.e. $(\boldsymbol{F}, \boldsymbol{Q})$ are valid instruments for $(\boldsymbol{F}, \boldsymbol{Q})$ themselves in the sense of standard IV theory [1, 3]. Then:*

1. *The factor-level causal parameter $\boldsymbol{\kappa}_0$ is identified from the moment conditions*

$$\mathbb{E}[(\boldsymbol{F}, \boldsymbol{Q})^\top(\boldsymbol{y} - \boldsymbol{F}\boldsymbol{\kappa}_0 - \boldsymbol{Q}\boldsymbol{\gamma}_0)] = \mathbf{0}.$$

*Moreover, any consistent IV or GMM estimator $\widehat{\boldsymbol{\kappa}}$ of $\boldsymbol{\kappa}_0$ satisfies $\widehat{\boldsymbol{\kappa}} \xrightarrow{p} \boldsymbol{\kappa}_0$ as $n \to \infty$.*

2. *The per-feature causal effect* $\boldsymbol{\alpha}_0 = \boldsymbol{V}\boldsymbol{\kappa}_0$ *satisfies the IV moment conditions*

$$\mathbb{E}[\boldsymbol{F}^\top(\boldsymbol{y} - \boldsymbol{X}\boldsymbol{\alpha}_0 - \boldsymbol{Q}\boldsymbol{\phi}_0)] = \mathbf{0}, \qquad \mathbb{E}[\boldsymbol{Q}^\top(\boldsymbol{y} - \boldsymbol{X}\boldsymbol{\alpha}_0 - \boldsymbol{Q}\boldsymbol{\phi}_0)] = \mathbf{0}, \tag{8}$$

*and is thus identified as the unique solution to these moment equations under the rank condition (IV4).*

*Proof.* We proceed in two steps.

**Step 1: Factor-level identification.** Equation (7) can be written as

$$\mathbf{y} = \mathbf{W}\boldsymbol{\theta}_0 + \mathbf{u}, \quad \mathbf{W} = (\mathbf{F}, \mathbf{Q}), \quad \boldsymbol{\theta}_0 = \begin{pmatrix} \boldsymbol{\kappa}_0 \\ \gamma_0 \end{pmatrix}.$$

The IV moment conditions are

$$\mathbb{E}[\mathbf{W}^\top \mathbf{u}] = \mathbf{0}.$$

Under (IV1)–(IV4), the standard theory of linear IV and GMM [1, 3, 6] implies that $\boldsymbol{\theta}_0$ is identified as the unique vector satisfying

$$\mathbb{E}[\mathbf{W}^\top(\mathbf{y} - \mathbf{W}\boldsymbol{\theta}_0)] = \mathbf{0},$$

provided that $\mathbb{E}[\mathbf{W}^\top\mathbf{W}]$ has full column rank. This yields identification of $\boldsymbol{\kappa}_0$ and, under mild regularity conditions (e.g. finite fourth moments, non-singular instrument covariance), consistency of the 2SLS or GMM estimator $\widehat{\boldsymbol{\kappa}}$ for $\boldsymbol{\kappa}_0$.

**Step 2: Per-feature causal effects and moment conditions.** We now relate the per-feature causal parameter $\boldsymbol{\alpha}_0$ to the factor-level parameter $\boldsymbol{\kappa}_0$. From (4) and the definition $\boldsymbol{\kappa}_0 = \mathbf{V}^\top\boldsymbol{\alpha}_0$, we have

$$\boldsymbol{\alpha}_0 = \mathbf{V}\boldsymbol{\kappa}_0.$$

Substituting the first-stage factor model (6) into the structural outcome model (1), we can rewrite

$$\mathbf{y} = \mathbf{X}\boldsymbol{\alpha}_0 + \mathbf{Q}\boldsymbol{\phi}_0 + \boldsymbol{\epsilon} = (\mathbf{F}\mathbf{V}^\top + \mathbf{Q}\boldsymbol{\psi} + \mathbf{E})\boldsymbol{\alpha}_0 + \mathbf{Q}\boldsymbol{\phi}_0 + \boldsymbol{\epsilon}.$$

Using $\boldsymbol{\alpha}_0 = \mathbf{V}\boldsymbol{\kappa}_0$, this becomes

$$\begin{aligned} \mathbf{y} &= \mathbf{F}\mathbf{V}^\top\mathbf{V}\boldsymbol{\kappa}_0 + \mathbf{Q}\boldsymbol{\psi}\mathbf{V}\boldsymbol{\kappa}_0 + \mathbf{E}\mathbf{V}\boldsymbol{\kappa}_0 + \mathbf{Q}\boldsymbol{\phi}_0 + \boldsymbol{\epsilon} \\ &= \mathbf{F}\boldsymbol{\kappa}_0 + \mathbf{Q}(\boldsymbol{\psi}\mathbf{V}\boldsymbol{\kappa}_0 + \boldsymbol{\phi}_0) + (\mathbf{E}\mathbf{V}\boldsymbol{\kappa}_0 + \boldsymbol{\epsilon}), \end{aligned}$$

where we again identify

$$\gamma_0 = \boldsymbol{\psi}\mathbf{V}\boldsymbol{\kappa}_0 + \boldsymbol{\phi}_0, \qquad \mathbf{u} = \mathbf{E}\mathbf{V}\boldsymbol{\kappa}_0 + \boldsymbol{\epsilon}.$$

Thus, (7) is recovered, and the factor-level moment conditions

$$\mathbb{E}[\mathbf{F}^\top(\mathbf{y} - \mathbf{F}\boldsymbol{\kappa}_0 - \mathbf{Q}\gamma_0)] = \mathbf{0}, \qquad \mathbb{E}[\mathbf{Q}^\top(\mathbf{y} - \mathbf{F}\boldsymbol{\kappa}_0 - \mathbf{Q}\gamma_0)] = \mathbf{0}$$

hold by assumption.

On the other hand, from (1), we can write

$$\mathbf{y} - \mathbf{X}\boldsymbol{\alpha}_0 - \mathbf{Q}\boldsymbol{\phi}_0 = \boldsymbol{\epsilon}.$$

Thus,

$$\mathbb{E}[\mathbf{F}^\top(\mathbf{y} - \mathbf{X}\boldsymbol{\alpha}_0 - \mathbf{Q}\boldsymbol{\phi}_0)] = \mathbb{E}[\mathbf{F}^\top\boldsymbol{\epsilon}] = \mathbf{0},$$

by Theorem 1. Likewise, since $\mathbf{Q}$ is included as an exogenous regressor and is assumed orthogonal to $\boldsymbol{\epsilon}$ in the usual way,

$$\mathbb{E}[\mathbf{Q}^\top(\mathbf{y} - \mathbf{X}\boldsymbol{\alpha}_0 - \mathbf{Q}\boldsymbol{\phi}_0)] = \mathbb{E}[\mathbf{Q}^\top\boldsymbol{\epsilon}] = \mathbf{0}.$$

This establishes the moment conditions (8) and shows that $\boldsymbol{\alpha}_0 = \mathbf{V}\boldsymbol{\kappa}_0$ is the unique solution under the rank condition (IV4). Standard arguments from linear IV and GMM theory [3, 6] then imply consistency of the corresponding estimator $\widehat{\boldsymbol{\alpha}} = \widehat{\mathbf{V}}\,\widehat{\boldsymbol{\kappa}}$ when $(\mathbf{U}, \mathbf{D}, \mathbf{V})$ are consistently estimated in the first stage. □

### S5. Exogeneity of Instrumental Factors in the Generalized First-Stage Model

In this section we show that the instrumental latent factors $\mathbf{F} = \mathbf{ZUD}$ are valid instruments in the generalized linear first-stage model used for high-dimensional count data (e.g. microbiome sequencing), working entirely in the observed-data space. The result extends the exogeneity argument from the Gaussian case to negative binomial (NB) and Poisson likelihoods commonly used for sequencing counts, under standard IV assumptions on the observed variables [1, 6, 3].

**Lemma 1** (Exogeneity of Factor IV instrumental factors in the generalized NB/Poisson first-stage model)**.** *Let $(\boldsymbol{Z}, \boldsymbol{Q}, \boldsymbol{X}, \boldsymbol{y})$ denote the observed data, where $\boldsymbol{Z} \in \mathbb{R}^{n\times d}$ are instruments, $\boldsymbol{Q} \in \mathbb{R}^{n\times t}$ are observed confounders, $\boldsymbol{X} \in \mathbb{R}^{n\times p}$ are endogenous features (e.g. microbiome counts), and $\boldsymbol{y} \in \mathbb{R}^n$ is the outcome. Assume:*

1. ***Outcome model (observed-data structural equation).*** *The outcome satisfies a generalized linear structural model*

$$h(\mathbf{E}[y_i \mid \boldsymbol{X}_i, \boldsymbol{Q}_i]) \;=\; \boldsymbol{X}_i^\top\boldsymbol{\alpha}_0 + \boldsymbol{Q}_i^\top\boldsymbol{\phi}_0, \qquad i = 1, \dots, n, \tag{9}$$

   *for some link function $h(\cdot)$, with structural errors $\epsilon_i = y_i - \mathbf{E}[y_i \mid \boldsymbol{X}_i, \boldsymbol{Q}_i]$ collected in $\boldsymbol{\epsilon} = (\epsilon_1, \dots, \epsilon_n)^\top$. By construction, $\mathbf{E}[\boldsymbol{\epsilon} \mid \boldsymbol{X}, \boldsymbol{Q}] = \mathbf{0}$.*

2. ***Generalized first-stage model for counts.*** *For each $i = 1, \dots, n$ and $j = 1, \dots, p$,*

$$X_{ij} \mid \boldsymbol{Z}_i, \boldsymbol{Q}_i \sim \mathrm{NB}(\mu_{ij}, \varphi_j), \qquad \log\mu_{ij} = \eta_{ij} = \boldsymbol{Z}_i^\top\beta_j + \boldsymbol{Q}_i^\top\psi_j, \tag{10}$$

   *where $\varphi_j > 0$ is a dispersion parameter and $\beta_j \in \boldsymbol{R}^d$, $\psi_j \in \boldsymbol{R}^t$ are coefficient vectors. Let $\boldsymbol{B} = (\beta_1, \dots, \beta_p) \in \boldsymbol{R}^{d\times p}$ and $\Psi = (\psi_1, \dots, \psi_p) \in \boldsymbol{R}^{t\times p}$.*

3. ***Low-rank structure in the instrument–feature map.*** *The instrument–feature coefficient matrix admits a sparse low-rank factorization*

$$\boldsymbol{B} = \boldsymbol{UDV}^\top, \qquad \boldsymbol{U} \in \boldsymbol{R}^{d\times r},\; \boldsymbol{D} = \mathrm{diag}(d_1, \dots, d_r),\; \boldsymbol{V} \in \boldsymbol{R}^{p\times r}, \quad r \ll \min\{d, p\}, \tag{11}$$

   *and we define the* instrumental factors

$$\boldsymbol{F} = \boldsymbol{ZUD} \in \boldsymbol{R}^{n\times r}. \tag{12}$$

4. ***Observed-data IV assumptions.***

   (a) ***Relevance:*** $\operatorname{rank}(\boldsymbol{B}) = r$ *and* $\operatorname{Cov}(\boldsymbol{Z}, \boldsymbol{X} \mid \boldsymbol{Q}) \neq 0$*, i.e. the instruments explain variation in* $\boldsymbol{X}$ *beyond* $\boldsymbol{Q}$*.*

   (b) ***Exogeneity:*** *the instruments are independent of the structural error given observed confounders,*

$$\boldsymbol{Z} \perp\!\!\!\perp \boldsymbol{\epsilon} \,\Big|\, \boldsymbol{Q}. \tag{13}$$

   *This is the standard IV exogeneity condition in the observed-data model [3].*

   (c) ***Exclusion restriction:*** *conditional on* $(\boldsymbol{X}, \boldsymbol{Q})$*, the instruments have no direct effect on* $\boldsymbol{y}$*,*

$$\boldsymbol{y} \mid \boldsymbol{X}, \boldsymbol{Q}, \boldsymbol{Z} \overset{d}{=} \boldsymbol{y} \mid \boldsymbol{X}, \boldsymbol{Q}. \tag{14}$$

*Then the instrumental factors are conditionally exogenous:*

$$\operatorname{Cov}(\boldsymbol{F}, \boldsymbol{\epsilon} \mid \boldsymbol{Q}) \;=\; \mathbf{0}, \tag{15}$$

*so that* $\boldsymbol{F}$ *can be used as a valid low-dimensional instrument set for identifying the causal parameter* $\boldsymbol{\alpha}_0$*.*

*Proof outline.* The argument is a direct consequence of (1)–(4) and the fact that $\mathbf{F}$ is a measurable function of the instruments $\mathbf{Z}$.

**Step 1: F is a deterministic function of Z.** By definition,

$$\mathbf{F} = \mathbf{ZUD}.$$

Given $(\mathbf{U}, \mathbf{D})$, the matrix $\mathbf{F}$ is fully determined by $\mathbf{Z}$, so $\mathbf{F}$ is measurable with respect to the $\sigma$-field generated by $(\mathbf{Z}, \mathbf{Q})$. Any randomness in $\mathbf{F}$ arises solely through $\mathbf{Z}$ (for fixed $\mathbf{U}, \mathbf{D}$ obtained from the first-stage fit).

**Step 2: Exogeneity of Z transfers to F.** Assumption (13) states that

$$\mathbf{Z} \perp\!\!\!\perp \boldsymbol{\epsilon} \,\Big|\, \mathbf{Q}.$$

If $g(\cdot)$ is any measurable function, then $g(\mathbf{Z})$ is also conditionally independent of $\boldsymbol{\epsilon}$ given $\mathbf{Q}$. Choosing $g(\mathbf{Z}) = \mathbf{ZUD}$ yields

$$\mathbf{F} = g(\mathbf{Z}) \perp\!\!\!\perp \boldsymbol{\epsilon} \,\Big|\, \mathbf{Q}.$$

**Step 3: Zero conditional covariance.** Using the law of iterated expectations and the measurability of $\mathbf{F}$ with respect to $(\mathbf{Z}, \mathbf{Q})$, we obtain

$$\begin{aligned}
\mathbb{E}[\mathbf{F}^\top \boldsymbol{\epsilon} \mid \mathbf{Q}] &= \mathbb{E}\Big[\mathbb{E}[\mathbf{F}^\top \boldsymbol{\epsilon} \mid \mathbf{Z}, \mathbf{Q}] \,\Big|\, \mathbf{Q}\Big] \\
&= \mathbb{E}\Big[\mathbf{F}^\top \mathbb{E}[\boldsymbol{\epsilon} \mid \mathbf{Z}, \mathbf{Q}] \,\Big|\, \mathbf{Q}\Big].
\end{aligned}$$

By conditional exogeneity (13), $\mathbb{E}[\boldsymbol{\epsilon} \mid \mathbf{Z}, \mathbf{Q}] = \mathbb{E}[\boldsymbol{\epsilon} \mid \mathbf{Q}]$. Moreover, by the definition of the structural error in (9), $\mathbb{E}[\boldsymbol{\epsilon} \mid \mathbf{X}, \mathbf{Q}] = \mathbf{0}$, and hence in particular $\mathbb{E}[\boldsymbol{\epsilon} \mid \mathbf{Q}] = \mathbf{0}$. Thus

$$\mathbb{E}[\boldsymbol{\epsilon} \mid \mathbf{Z}, \mathbf{Q}] = \mathbf{0},$$

and therefore

$$\mathbb{E}[\mathbf{F}^\top \boldsymbol{\epsilon} \mid \mathbf{Q}] = \mathbb{E}[\mathbf{F}^\top \cdot \mathbf{0} \mid \mathbf{Q}] = \mathbf{0}.$$

This implies $\mathrm{Cov}(\mathbf{F}, \boldsymbol{\epsilon} \mid \mathbf{Q}) = \mathbf{0}$.

**Step 4: Role of the NB/Poisson first stage.** The negative binomial (or Poisson) first-stage model (10) and the low-rank structure (11) are used to construct a low-dimensional, instrument-induced summary $\mathbf{F}$ and to ensure relevance (assumption (4)(a)). The exogeneity result itself does not depend on the specific likelihood for $\mathbf{X}$, but only on the observed-data IV assumptions (13)–(14) and on the fact that $\mathbf{F}$ is a deterministic transformation of $\mathbf{Z}$ [1, 6, 3]. □

### S6. Improved Causal Identification under Multi-Omics Factor IV

We formalize how expanding the endogenous measurement space from a single-omics matrix $\mathbf{X}$ to a multi-omics matrix $\mathbf{X}'$ strengthens the plausibility of the exclusion restriction in the Factor IV framework, thereby improving identification of the causal effect of instrument-aligned latent factors on the outcome.

We consider the observed-data model used throughout the paper. Let $\mathbf{Z} \in \mathbb{R}^{n\times d}$ denote instrumental variables, $\mathbf{Q} \in \mathbb{R}^{n\times t}$ observed confounders, $\mathbf{X} \in \mathbb{R}^{n\times p}$ endogenous features, and $\mathbf{y} \in \mathbb{R}^n$ the outcome. Factor IV constructs latent instrumental factors $\mathbf{F} \in \mathbb{R}^{n\times r}$ as deterministic functions of $\mathbf{Z}$ via a low-rank instrument–exposure map, and estimates causal effects through the second-stage model $\mathbf{y} = \mathbf{F}\boldsymbol{\kappa} + \mathbf{Q}\boldsymbol{\phi} + \boldsymbol{\epsilon}$.

We distinguish between:

- **Single-omics setting:** $\mathbf{X}$ contains features from a single molecular layer (e.g., microbiome).
- **Multi-omics setting:** $\mathbf{X}' = (\mathbf{X}, \mathbf{M})$ augments $\mathbf{X}$ with additional molecular layers $\mathbf{M}$ (e.g., metabolites, transcripts), where $\dim(\mathbf{X}') > \dim(\mathbf{X})$.

Here, $\mathbf{M}$ denotes a latent mediator affected by $\mathbf{Z}$ that lies on a causal path to $\mathbf{y}$ but is unobserved in the single-omics analysis. The core identifying assumption is the exclusion restriction for the latent factors:

$$\mathbf{y} \perp\!\!\!\perp \mathbf{Z} \mid \mathbf{F}, \mathbf{Q}, \tag{16}$$

meaning that instruments affect the outcome only through the factor-aligned endogenous variation.

##### Exclusion violation under single-omics measurements

In the single-omics setting, the latent factors $\mathbf{F}^{\text{single}}$ are constructed from $\mathbf{Z}$ and $\mathbf{X}$ alone. Suppose there exists an unmeasured mediator $\mathbf{M}$ such that

$$\mathbf{Z} \longrightarrow \mathbf{M} \longrightarrow \mathbf{y},$$

with $\mathbf{M} \not\subset \mathbf{X}$. Because $\mathbf{M}$ is omitted, the factor construction step cannot capture the full $\mathbf{Z}$-induced variation that influences $\mathbf{y}$. As a result, conditioning on $\mathbf{F}^{\text{single}}$ fails to block the path $\mathbf{Z} \to \mathbf{M} \to \mathbf{y}$, yielding

$$\mathbf{y} \not\perp\!\!\!\perp \mathbf{Z} \mid \mathbf{F}^{\text{single}}, \mathbf{Q},$$

and bias in the estimated causal effect of $\mathbf{F}^{\text{single}}$.

#### Strengthening exclusion via multi-omics factorization

In the multi-omics setting, the endogenous space is expanded to $\mathbf{X}' = (\mathbf{X}, \mathbf{M})$, and latent factors $\mathbf{F}^{\text{multi}}$ are constructed from the joint instrument–exposure relationship. This expansion has two key consequences. First, because $\mathbf{M} \subset \mathbf{X}'$, any $\mathbf{Z}$-induced variation in $\mathbf{M}$ is necessarily represented in the factor space. The factors $\mathbf{F}^{\text{multi}}$ therefore act as a more complete proxy for the total set of mediators through which $\mathbf{Z}$ influences $\mathbf{y}$. Second, conditioning on $\mathbf{F}^{\text{multi}}$ blocks the previously unmeasured causal path:

$$\mathbf{Z} \longrightarrow \mathbf{X}' \text{ (including } \mathbf{M}) \longrightarrow \mathbf{F}^{\text{multi}} \longrightarrow \mathbf{y}.$$

In graphical terms, $\mathbf{F}^{\text{multi}}$ d-separates $\mathbf{Z}$ and $\mathbf{y}$ along paths mediated by $\mathbf{M}$, restoring the exclusion restriction:

$$\mathbf{y} \perp\!\!\!\perp \mathbf{Z} \mid \mathbf{F}^{\text{multi}}, \mathbf{Q}.$$

#### Implications for identification

By incorporating additional molecular layers that mediate instrument effects, the multi-omics Factor IV construction reduces violations of the exclusion restriction arising from unmeasured biological pathways. While exclusion cannot be empirically verified, expanding $\mathbf{X}$ to $\mathbf{X}'$ increases the plausibility of the identifying assumptions and yields factor-level causal effects $\boldsymbol{\kappa}$ and structured feature-level effects $\boldsymbol{\alpha} = \mathbf{V}\boldsymbol{\kappa}$ that are less susceptible to bias from omitted mediation. This provides a formal justification for improved causal identification under multi-omics integration within the Factor IV framework.

### S7. Additional Analysis of Figures

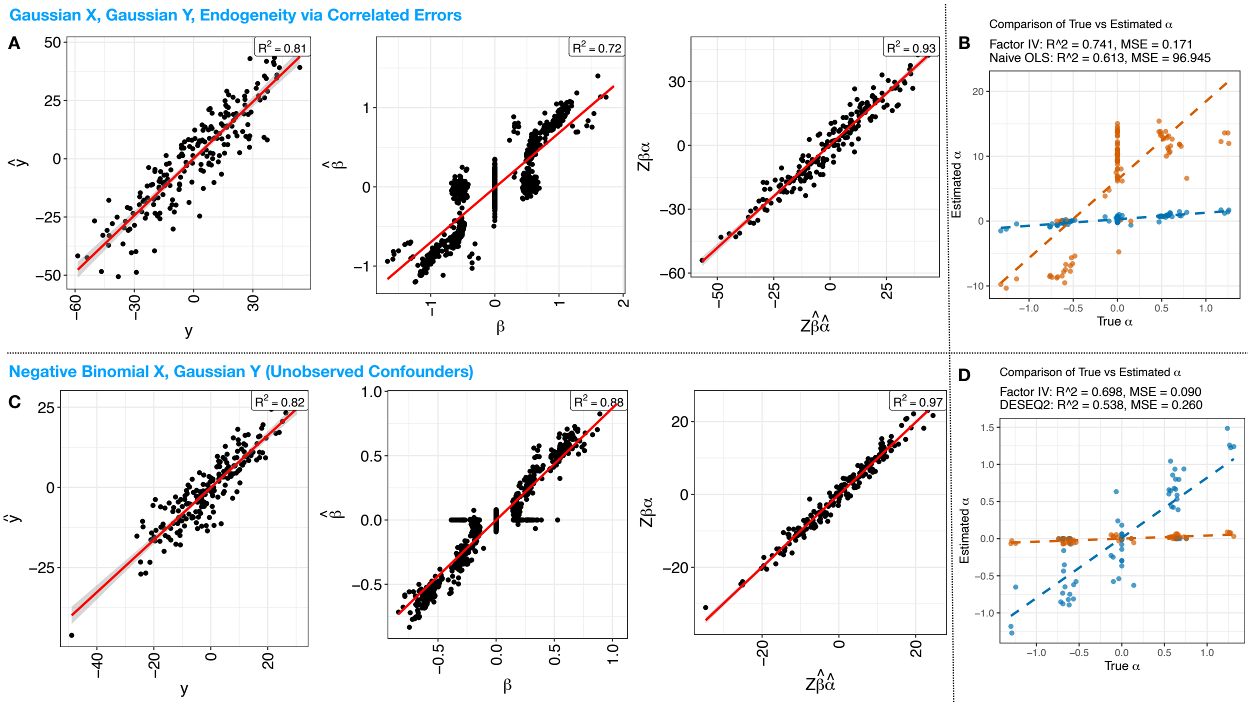


Figure 1: **Simulation results evaluating the Factor IV framework.** Simulation studies under Gaussian and negative binomial exposure models assessing recovery of causal structure in the presence of endogeneity arising from correlated errors and unobserved confounding, respectively. (**A**) Gaussian setting: comparison of observed and predicted outcomes $\mathbf{Y}$, instrument–exposure signal $\mathbf{Z}\boldsymbol{\beta}$, and the projected causal component $\mathbf{Z}\boldsymbol{\beta}\hat{\boldsymbol{\alpha}}$. (**B**) Gaussian setting: comparison of estimated feature-level causal effects $\hat{\boldsymbol{\alpha}}$ from Factor IV versus a naive regression approach. (**C**) Negative binomial setting: comparison of observed and predicted outcomes $\mathbf{Y}$, instrument–exposure signal $\mathbf{Z}\boldsymbol{\beta}$, and the projected causal component $\mathbf{Z}\boldsymbol{\beta}\hat{\boldsymbol{\alpha}}$. (**D**) Negative binomial setting: comparison of estimated causal effects $\hat{\boldsymbol{\alpha}}$ from Factor IV and DESeq2.

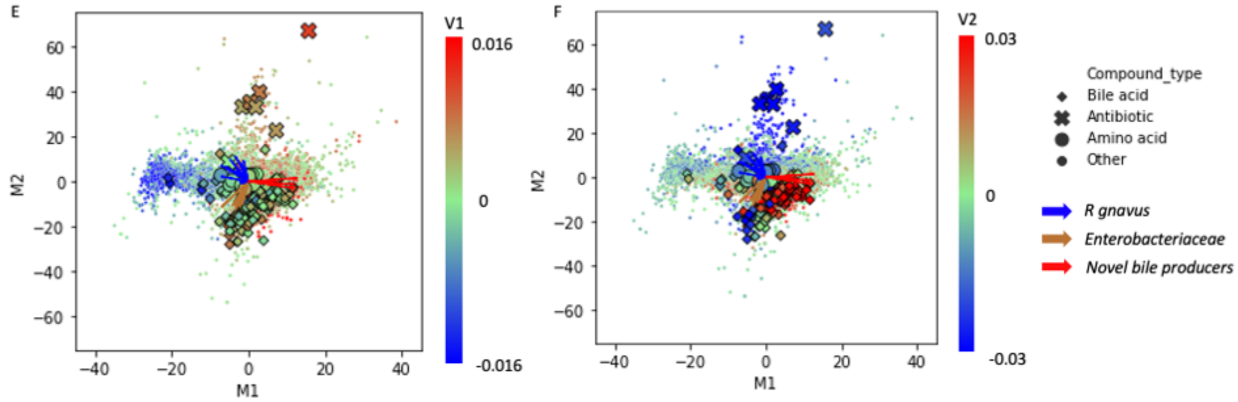


Figure 2: **Microbe–metabolite co-occurrence structure inferred by MMvec.** MMvec analysis [4] showing learned co-occurrence relationships between microbial taxa and metabolites from the HCC dataset. Points represent microbes and metabolites embedded in a shared latent space, with proximity indicating higher conditional co-occurrence. Major metabolite classes and representative taxa are annotated.

### S8. Equivalence Between an Unobserved Confounder and Correlated Errors

**Observed-data setup.** Throughout we work with the observed-data structural model $\mathbf{y} = \mathbf{X}\boldsymbol{\alpha}_0 + \mathbf{Q}\boldsymbol{\phi}_0 + \boldsymbol{\epsilon}$, and a first-stage model $\mathbf{X} = \mathbf{Z}\boldsymbol{\beta}_0 + \mathbf{Q}\boldsymbol{\psi}_0 + \mathbf{E}$, where endogeneity means $\mathrm{Cov}(\mathbf{X}, \boldsymbol{\epsilon} \mid \mathbf{Q}) \neq 0$, equivalently $\mathrm{Cov}(\mathbf{E}, \boldsymbol{\epsilon} \mid \mathbf{Q}) \neq 0$. We show that two common mechanisms for endogeneity—(i) an unobserved confounder and (ii) correlated errors—are algebraically equivalent representations of the same joint law for $(\mathbf{X}, \mathbf{y})$, under mild second-moment conditions.

#### A. Unobserved confounder implies correlated errors

**Assumption 1** (Latent-confounder representation)**.** *There exists a random vector $U \in \mathbb{R}^s$ (unobserved confounder) and random noises $(\widetilde{\mathbf{E}}, \widetilde{\boldsymbol{\epsilon}})$ such that*

$$\begin{aligned} \boldsymbol{X} &= \boldsymbol{Z}\boldsymbol{\beta}_0 + \boldsymbol{Q}\boldsymbol{\psi}_0 + U\Lambda_X^\top + \widetilde{\mathbf{E}}, \\ \boldsymbol{y} &= \boldsymbol{X}\boldsymbol{\alpha}_0 + \boldsymbol{Q}\boldsymbol{\phi}_0 + U\lambda_Y + \widetilde{\boldsymbol{\epsilon}}, \end{aligned}$$

*with $\mathbb{E}[U \mid \boldsymbol{Q}] = 0$, $\mathbb{E}[\widetilde{\mathbf{E}} \mid \boldsymbol{Q}] = 0$, $\mathbb{E}[\widetilde{\boldsymbol{\epsilon}} \mid \boldsymbol{Q}] = 0$, and $(\widetilde{\mathbf{E}}, \widetilde{\boldsymbol{\epsilon}}) \perp\!\!\!\perp U \mid \boldsymbol{Q}$. Here $\Lambda_X \in \mathbb{R}^{p \times s}$ and $\lambda_Y \in \mathbb{R}^s$ are loadings.*

**Lemma 2** (Latent confounding induces correlated errors)**.** *Under Assumption 1, define observed-data errors $\mathbf{E} := U\Lambda_X^\top + \widetilde{\mathbf{E}}$ and $\boldsymbol{\epsilon} := U\lambda_Y + \widetilde{\boldsymbol{\epsilon}}$. Then $\mathrm{Cov}(\mathbf{E}, \boldsymbol{\epsilon} \mid \boldsymbol{Q}) = \Lambda_X \Sigma_U \lambda_Y$, where $\Sigma_U := \mathrm{Cov}(U \mid \boldsymbol{Q})$, and hence $\mathrm{Cov}(\boldsymbol{X}, \boldsymbol{\epsilon} \mid \boldsymbol{Q}) \neq 0$ whenever $\Lambda_X \Sigma_U \lambda_Y \neq 0$.*

*Proof.* By definition and conditional mean-zero, $\mathrm{Cov}(\mathbf{E}, \boldsymbol{\epsilon} \mid \mathbf{Q}) = \mathrm{Cov}(U\Lambda_X^\top + \widetilde{\mathbf{E}}, U\lambda_Y + \widetilde{\boldsymbol{\epsilon}} \mid \mathbf{Q})$.

Expanding and using conditional independence of $(\widetilde{\mathbf{E}}, \widetilde{\boldsymbol{\epsilon}})$ from $U$ given $\mathbf{Q}$ yields

$$\begin{aligned}\operatorname{Cov}(\mathbf{E}, \boldsymbol{\epsilon} \mid \mathbf{Q}) &= \operatorname{Cov}(U\Lambda_X^\top,\, U\lambda_Y \mid \mathbf{Q}) + \operatorname{Cov}(\widetilde{\mathbf{E}}, \widetilde{\boldsymbol{\epsilon}} \mid \mathbf{Q}) \\ &= \Lambda_X \operatorname{Cov}(U \mid \mathbf{Q})\, \lambda_Y + \operatorname{Cov}(\widetilde{\mathbf{E}}, \widetilde{\boldsymbol{\epsilon}} \mid \mathbf{Q}).\end{aligned}$$

If we take $(\widetilde{\mathbf{E}}, \widetilde{\boldsymbol{\epsilon}})$ conditionally uncorrelated given $\mathbf{Q}$ (a standard normalization), the second term is 0 and the displayed identity follows. Since $\mathbf{X} = \mathbf{Z}\boldsymbol{\beta}_0 + \mathbf{Q}\boldsymbol{\psi}_0 + \mathbf{E}$ with $(\mathbf{Z}, \mathbf{Q})$ exogenous, $\operatorname{Cov}(\mathbf{X}, \boldsymbol{\epsilon} \mid \mathbf{Q}) = \operatorname{Cov}(\mathbf{E}, \boldsymbol{\epsilon} \mid \mathbf{Q})$. □

#### B. Correlated errors admit an equivalent latent-confounder representation

**Assumption 2** (Correlated-error representation)**.** *The observed-data errors $(\mathbf{E}, \boldsymbol{\epsilon})$ satisfy $\mathbb{E}[\mathbf{E} \mid \boldsymbol{Q}] = 0$, $\mathbb{E}[\boldsymbol{\epsilon} \mid \boldsymbol{Q}] = 0$ and have conditional covariance $\Sigma_{E\epsilon} := \operatorname{Cov}(\mathbf{E}, \boldsymbol{\epsilon} \mid \boldsymbol{Q}) \in \mathbb{R}^{p\times 1}$, with finite second moments. No parametric distribution is required.*

**Theorem 2** (Equivalence: correlated errors $\iff$ latent confounder)**.** *Under Assumption 2, there exists a latent variable $U \in \mathbb{R}^s$ (for some $s \le p+1$), loadings $(\Lambda_X, \lambda_Y)$, and residual noises $(\widetilde{\mathbf{E}}, \widetilde{\boldsymbol{\epsilon}})$ such that*

$$\begin{aligned}\mathbf{E} &= U\Lambda_X^\top + \widetilde{\mathbf{E}}, \\ \boldsymbol{\epsilon} &= U\lambda_Y + \widetilde{\boldsymbol{\epsilon}},\end{aligned}$$

*with $(\widetilde{\mathbf{E}}, \widetilde{\boldsymbol{\epsilon}}) \perp\!\!\!\perp U \mid \boldsymbol{Q}$ and $\operatorname{Cov}(\widetilde{\mathbf{E}}, \widetilde{\boldsymbol{\epsilon}} \mid \boldsymbol{Q}) = 0$, and moreover $\operatorname{Cov}(\mathbf{E}, \boldsymbol{\epsilon} \mid \boldsymbol{Q}) = \Lambda_X \Sigma_U \lambda_Y$, so the endogeneity induced by $\Sigma_{E\epsilon}$ can be represented as arising from an unobserved confounder. Conversely, any latent-confounder model of the form in Assumption 1 implies correlated errors as in Lemma 2. Thus the two representations are observationally equivalent at the level of second moments of $(\boldsymbol{X}, \boldsymbol{y})$ given $(\boldsymbol{Z}, \boldsymbol{Q})$.*

*Proof.* We give a constructive proof using a one-factor version ($s = 1$), which already captures any *vector* cross-covariance $\Sigma_{E\epsilon}$ by choosing appropriate loadings.

Let $U$ be a scalar with $\mathbb{E}[U \mid \mathbf{Q}] = 0$ and $\operatorname{Var}(U \mid \mathbf{Q}) = 1$ (e.g., $U \sim \mathcal{N}(0,1)$ independent of $(\mathbf{Z}, \mathbf{Q})$). Define loadings $\Lambda_X := \Sigma_{E\epsilon} \in \mathbb{R}^p$ and $\lambda_Y := 1$. Now define residuals $\widetilde{\boldsymbol{\epsilon}} := \boldsymbol{\epsilon} - U$ and $\widetilde{\mathbf{E}} := \mathbf{E} - U\Lambda_X^\top$. By construction, $\mathbf{E} = U\Lambda_X^\top + \widetilde{\mathbf{E}}$ and $\boldsymbol{\epsilon} = U\lambda_Y + \widetilde{\boldsymbol{\epsilon}}$.

Compute the cross-covariance:

$$\begin{aligned}\operatorname{Cov}(\mathbf{E}, \boldsymbol{\epsilon} \mid \mathbf{Q}) &= \operatorname{Cov}(U\Lambda_X^\top + \widetilde{\mathbf{E}},\, U + \widetilde{\boldsymbol{\epsilon}} \mid \mathbf{Q}) \\ &= \Lambda_X \operatorname{Var}(U \mid \mathbf{Q})\, 1 + \operatorname{Cov}(\widetilde{\mathbf{E}}, \widetilde{\boldsymbol{\epsilon}} \mid \mathbf{Q}) + \underbrace{\operatorname{Cov}(U\Lambda_X^\top, \widetilde{\boldsymbol{\epsilon}} \mid \mathbf{Q})}_{(*)} + \underbrace{\operatorname{Cov}(\widetilde{\mathbf{E}}, U \mid \mathbf{Q})}_{(**)}.\end{aligned}$$

We now *choose* $(\widetilde{\mathbf{E}}, \widetilde{\boldsymbol{\epsilon}})$ so that: (i) $\operatorname{Cov}(\widetilde{\mathbf{E}}, \widetilde{\boldsymbol{\epsilon}} \mid \mathbf{Q}) = 0$, (ii) $\operatorname{Cov}(U, \widetilde{\boldsymbol{\epsilon}} \mid \mathbf{Q}) = 0$, and (iii) $\operatorname{Cov}(U, \widetilde{\mathbf{E}} \mid \mathbf{Q}) = 0$. This is always possible by defining $U$ as the (conditional) linear projection component shared by $(\mathbf{E}, \boldsymbol{\epsilon})$. Concretely, in the Gaussian case this is exactly the standard decomposition of a jointly Gaussian vector into a common factor plus independent residuals, and in the general finite-variance case it follows from orthogonalization (Gram–Schmidt) in $L_2$.

Under (ii) and (iii), terms $(*)$ and $(**)$ vanish, and since $\operatorname{Var}(U \mid \mathbf{Q}) = 1$ we get $\operatorname{Cov}(\mathbf{E}, \boldsymbol{\epsilon} \mid \mathbf{Q}) = \Lambda_X = \Sigma_{E\epsilon}$, which matches the original correlated-error model. Hence the correlated-error structure can be represented via a latent confounder $U$. The converse direction is Lemma 2. Therefore, the two mechanisms are observationally equivalent (at least up to second moments). □

**Remarks (interpretation for simulation design).** Theorem 2 explains why simulation mechanisms based on (i) an explicit latent confounder entering both stages and (ii) a shared noise component inducing $\mathrm{Cov}(\mathbf{E}, \boldsymbol{\epsilon}) \neq 0$ are interchangeable: both generate the same kind of endogeneity in the observed-data model. In practice, the two parameterizations differ mainly in *interpretability* (biological confounding versus shared technical noise) rather than statistical consequences for IV identification.

**Connection to IV identification.** Because both mechanisms only affect the model through $\mathrm{Cov}(\mathbf{X}, \boldsymbol{\epsilon} \mid \mathbf{Q}) \neq 0$, valid instruments $\mathbf{Z}$ satisfying relevance and conditional exogeneity ($\mathbf{Z} \perp\!\!\!\perp \boldsymbol{\epsilon} \mid \mathbf{Q}$) remain the route to identification in either case [1, 3, 6].
